# The ecdysone receptor promotes or suppresses proliferation according to ligand level

**DOI:** 10.1101/2023.02.10.527985

**Authors:** Gantas Perez-Mockus, Luca Cocconi, Cyrille Alexandre, Birgit Aerne, Guillaume Salbreux, Jean-Paul Vincent

## Abstract

Steroid hormones control various cellular activities in a context-dependent manner. For example, ecdysone, which acts through a type II nuclear receptor, has seemingly opposite effects in Drosophila wing precursors, promoting proliferation during larval stages, and triggering proliferation arrest at pupariation. We find that wing precursors proliferate normally in the complete absence of the ecdysone receptor (EcR), whether ecdysone is present or not, suggesting that ecdysone overrides a default antiproliferative activity of the receptor. By contrast, termination of proliferation by high concentration of 20E at the end of larval life involves conventional gene regulation by the ligand-receptor complex. The switch from one mode of regulation to the other is determined by ligand level, as measured with a calibrated EcR transcriptional reporter and *ex vivo* proliferation assays. Accordingly, RNA Seq analysis uncovers distinct transcriptional responses to different doses of ecdysone. Some genes are only activated at high doses (high threshold targets) and likely to comprise genes that stop proliferation at pupariation, when ecdysone titres are high. We find that other target genes respond to all physiological concentrations of ecdysone. Some of these genes are known to promote proliferation and could therefore contribute to the pro-proliferation activity of low-level ecdysone. Finally, we show mathematically and with synthetic reporters that relatively simple combinations of regulatory elements can recapitulate the behaviour of both types of target genes.

## INTRODUCTION

Type II nuclear receptors constitute a subclass of nuclear receptors, transcription factors that bind small lipophilic molecules and mediate their signalling activity during development and adult homeostasis. One example is the retinoic acid receptor, which regulates germ layer formation, body axis formation, neurogenesis, cardiogenesis, and many other processes, during vertebrate development ^1-3^. Disruption of retinoic acid receptor signalling underlies numerous malignancies in adults and children. Another example is the thyroid hormone receptor, an important component of the endocrine system, which leads to a wide range of symptoms when misregulated ^4,5^. Understanding how type II nuclear receptors control these myriad functions is therefore essential. While type I receptors are regulated by ligand-induced nuclear import, type II receptors remain bound to DNA regardless of the ligand binding status, usually as heterodimers (e.g. with the retinoid X receptor) ^6^. In the absence of ligand, these heterodimers recruit a corepressor, which is replaced by a co-activator upon ligand binding. Thus, type II nuclear receptors can act either as transcription repressor or activator. It is generally thought that they regulate a wide range of activities by interacting with tissue-or stage-specific co-factors ^7^. In some case, type II receptors seem to have opposite effects on the same process, e.g. stimulating or suppressing proliferation depending on the context ^8-10^. The molecular basis of this feature remains poorly understood.

The main type II nuclear receptor of Drosophila is the ecdysone receptor (EcR). The EcR binds to DNA as a heterodimer with Ultraspiracle (USP) and, in the absence of ligand, recruits transcriptional corepressors such as Smrter ^11-15^ (Fig. 1 A). Ligand binding induces a conformational change that allows the recruitment of transcriptional co-activators ^12,16-19^. Production of the active hormone starts in the prothoracic gland (PG) through the concerted activity of enzymes encoded by the so-called halloween genes, which are themselves regulated by a combination of developmental, nutritional and stress signals (reviewed in ^20-24^). These enzymes transform cholesterol into ecdysone (E), which is secreted from the prothoracic gland to reach the larval fat body and gut where the hydroxyl-transferase encoded by *shade* ^25^ converts it into the active form, 20-hydroxy-ecdysone (20E). 20E is then released in the circulation and gains access to target tissues through a dedicated transporter ^26^.

**Figure 1:**
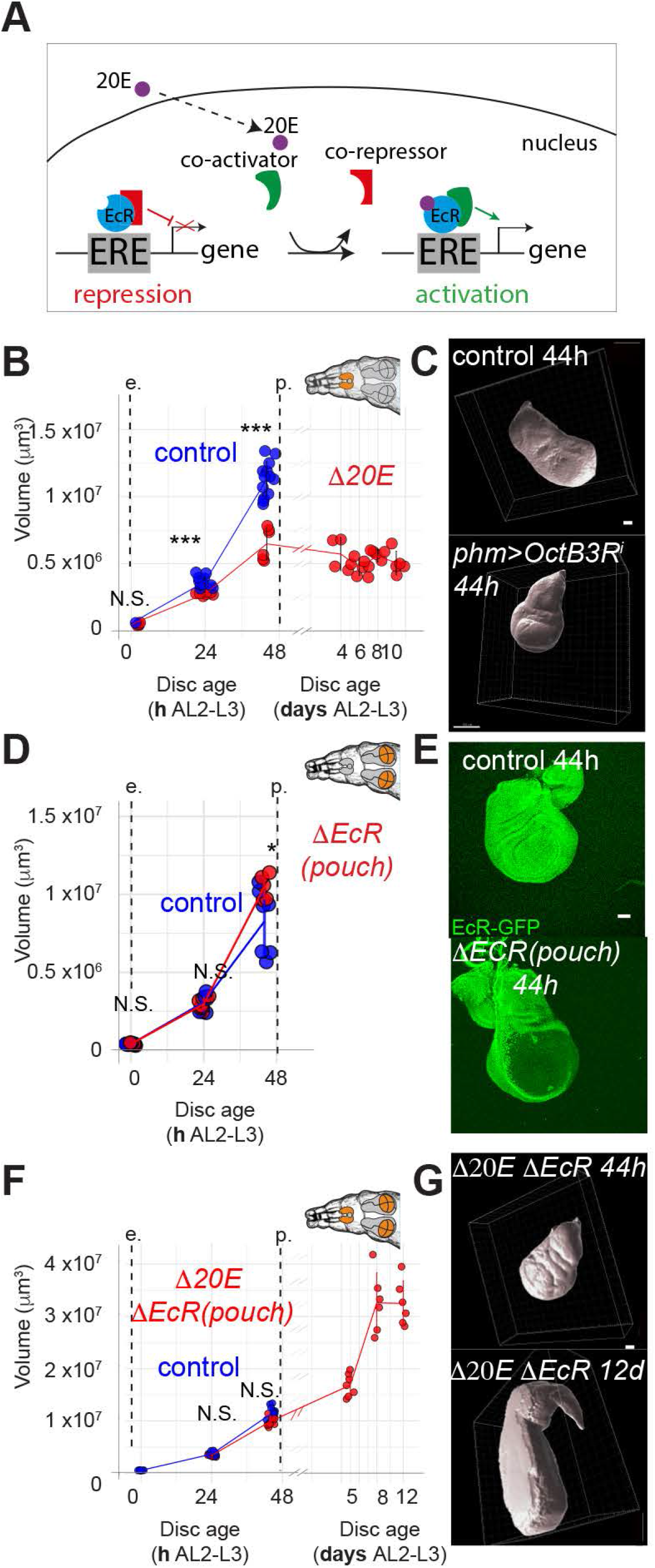
20E and EcR have opposite effects on wing disc growth. **A**. Schematic representation of EcR activity as a function of ligand availability. EcR is a transcriptional repressor in the absence of 20 (left) and becomes an activator with 20E (right). **B-C**. Inhibition of 20E synthesis (OctΔ3R knockdown in the prothoracic gland: phm> OctΔ3R^i^), impairs wing disc growth (quantification of volumetric reconstruction and representative images are shown, as in D-E and F-G below). **D-E**. Pouch-specific inactivation of the EcR (EcR_KO_ / EcR_cKO_ pdm2-Gal4 UAS-Flp) has no significant impact on disc growth. **F-G**. Pouch-specific inactivation of the EcR allows growth even when 20E synthesis is inhibited (EcR_KO_ LexOP-Flp/ EcR-GFP_cKO_; rotund-LexA UAS-OctΔ3R_RNAi_/ phm-gal4). In the absence of 20E, pupariation does not take place, allowing sustained disc growth beyond the normal time of pupariation. All error bars represent standard deviation. * p<0.5, *** p<0.01, N.S. No statistical difference. Wilcoxon Ranks sum tests were performed in B, D and F. Scale bars represent 50 μm.

Depletion of systemic 20E by genetic manipulation of the prothoracic gland leads to a marked slowdown of proliferation in wing precursors ^27-32^, suggesting that 20E is a proliferative signal. However, at the onset of metamorphosis, 20E leads to G2 arrest, in preparation for the morphogenetic rearrangements that subsequently take place ^33,34^. Therefore, 20E seems to promote proliferation during larval stages and prevent proliferation at pupariation.

How can the same molecular signal drive opposite effects on proliferation? To address this question, we devised new genetic tools to revisit the role of 20E and its receptor in wing imaginal disc proliferation. We found that, unexpectedly, in the absence of both receptor and ligand, proliferation proceeded normally. This leads us to propose that, prior to metamorphosis, 20E promotes growth by preventing the EcR from inhibiting growth/proliferation, a possibility raised previously by Schubiger and others ^35,36^. We devised a sensitive reporter of EcR signaling to estimate the range of 20E levels present at different stages and found that the low levels normally present *in vivo* stimulate proliferation *ex vivo*, while high levels are inhibitory. Bulk RNASeq of imaginal discs exposed to different concentration of 20E reveals that some target genes are only activated at high levels, while others respond to both low and high concentrations. Mathematical modelling, validated by synthetic reporters, shows that relatively simple changes in the cis-regulatory region of target genes could account for the qualitatively distinct responses of 20E target genes. We suggest that proliferation is actively suppressed by one or several high threshold genes, yet to be identified. Genes that promote proliferation are likely to be found among those that expressed at low 20E doses. A molecular understanding of proliferation control by 20E remains elusive but our results suggest that high threshold anti-proliferation dominantly suppress the function of the myriad genes that promote proliferation.

## RESULTS

### Ecdysone is not required for tissue growth if the EcR is genetically ablated

As a prelude to assessing the role of 20E and its receptor in growth and proliferation, we measured the growth of wild type Drosophila wing imaginal discs during the third instar, using volume as a proxy for biomass ^37^. To accurately stage imaginal discs, larvae were selected at the L2-L3 transition, a well-defined developmental milestone, then allowed to grow for specific periods of time before the wing discs were dissected, fixed, stained with DAPI, and mounted in a drop of agar-containing a clearing agent. The volume was then calculated from 3D reconstructed confocal stacks ^38^ (Sup. Fig. 1A and B). This analysis revealed that, during the 48 hours of the 3^rd^ instar, wild type wing imaginal disc volume increases by about 27-fold, which agrees with previous reports suggesting that, during this time, imaginal discs cells undergo approximately 9 to 10 divisions ^39-43^. We cannot be sure that growth terminates at pupariation since extensive morphogenesis occurring at this time makes volume measurements difficult. However, it is clear that, at the end of the third instar, only occasional cells undergo mitosis, as assayed with anti-pH3 staining ^34,44^, suggesting that proliferation grinds to a halt at the onset of pupariation.

Using the above assay, we then re-assessed the effect of reducing systemic ecdysone levels on wing imaginal disc growth and proliferation during the 3^rd^ instar. Ecdysone is produced by the prothoracic gland in response to activation of the β3-octopamine receptor ^45^. It is then oxidized in peripheral tissue to 20-Hydroxyecdysone (20E), the active hormone. As previously shown ^29,45-47^, larvae expressing an RNAi transgene against the β3-octopamine receptor specifically in the prothoracic gland failed to metamorphose (Sup. Fig. 1C), confirming that 20E is essential for this developmental transition to take place. The wing imaginal discs within these animals grew poorly, especially during the second half of the third instar. They also failed to gain volume during the subsequent 8 days of extended larval period (Fig. 1B and C). Note, however, that this genetic manipulation does not completely abrogate ecdysone production since these animals do progress through earlier instars, which also require 20E. Nevertheless the observation that growth is markedly reduced during the third instar confirms and extends the earlier suggestion that 20E is essential for imaginal discs growth ^28-32^.

The requirement of 20E for imaginal discs growth seems at odds with other reports that RNAi-mediated knockdown of the 20E receptor specifically in imaginal discs does not to impair growth ^35,36,48^. Since RNAi may leave residual gene activity, we engineered the *EcR* locus so that it could be completely inactivated by Flp recombinase in a tissue-specific manner (Sup. Fig. 1D). Specifically, a DNA fragment encoding GFP was inserted at the 3’ end of the coding region and FRT sites were added to allow Flp-mediated excision of the last 4 exons and the additional GFP-coding sequences. The product of this allele, termed EcR-GFP^cKO^ was found in the nucleus (Sup. Fig. 1F), where the endogenous EcR is known to reside ^49^. Moreover, EcR-GFP^cKO^ homozygous flies showed no morphological defects and developed at the same rate as control, wild type larvae (Sup. Fig. 1E). We conclude therefore that, in the absence of Flp, this allele is fully functional. We then used *pdm2-Gal4* with, *UAS-Flp* to inactivate this allele in the prospective wing of hemizygous *EcR* larvae (EcR-GFP^cKO^/EcR^KO^; pdm2-Gal4 UAS-Flp). This had no effect either on developmental timing of the whole larva (Sup. Fig. 1G) nor or on the rate of imaginal disc growth; third instar imaginal discs lacking EcR grew at the same rate as wild type disc (Fig.1 D-E). We therefore conclude that the EcR is not required for imaginal disc growth, even though its ligand is.

One could explain the requirement of 20E, but not that of EcR, for imaginal disc growth by invoking the existence of a distinct receptor through which 20E would control growth. To assess this possibility, we created larvae that are impaired in Ecdysone production (phm-gal4 UAS-OctΔ3R^RNAi^) while at the same time lacking, from the time of the L2-L3 transition, the EcR in the wing pouch (rotund-LexA LexOP-Flp EcR-GFP^cKO^/EcR^KO^), named hereafter Δ20E^larva^ ΔEcR^pouch^. In contrast to the situation with OctΔ3R downregulation alone (ΔEcR^pouch^), wing imaginal discs from Δ20E^larva^ ΔEcR^pouch^ animals grew seemingly normally during the usual growth period, showing that 20E is not absolutely required for growth and hence that an alternative receptor is not involved. As expected from the loss of 20E, Δ20E^larva^ ΔEcR^pouch^ larvae failed to pupariate, providing potential extra time for growth. This allowed imaginal discs to reach 3 times the size of wild type discs 10 days after the onset of 3^rd^ instar (Fig. 1F-G). Thus, EcR is required for proliferation arrest in response to the pulse of 20E at pupariation. Accordingly, imaginal discs expressing an EcR RNAi transgene at the time of disc specification (with vg-Gal4 UAS-Flp Act5c-FRT-STOP-FRT-Gal4) within larvae producing ecdysone normally, overgrew somewhat beyond the time of pupariation (Sup. Fig. 1 H-J), perhaps because they cannot sense the growth terminating pulse of ecdysone. These observations suggest that, in the absence of 20E, EcR could act as a proliferation brake. During the growth period, 20E would release this brake, thus promoting proliferation. In contrast, at the end of the third instar, 20E would activate the EcR to trigger the morphogenetic anti-proliferation programme of pupariation ^33,50,51^. How could the same ligand have opposite effects on proliferation? One possibility is that low level 20E promotes growth and proliferation during the third instar, while the high levels present at pupariation would trigger proliferation arrest and morphogenesis.

### EcR activity rises during the third instar

Liquid chromatography–mass spectrometry (LC-MS) measurements suggest that imaginal discs are exposed to relatively low 20E levels during the third instar compared to pupariation ^52^. To estimate to what extent systemic levels of 20E translate in EcR signaling activity within imaginal discs, we devised a reporter comprising 5 alternating copies of two consensus EcR response element (ERE) ^49,53^, upstream of a minimal heat shock promoter driving transcription of DNA encoding nuclear-targeted neonGreen tetramers (ERE-NLS4xNG) (Fig. 2A). Nuclear fluorescence was readily detected in transgenic imaginal discs at the end of the 3^rd^ instar. This signal was abrogated by expression of a dominant negative form of EcR (Sup. Fig. 2A). Fluorescence was also detected in live pupae, in a temporal pattern that mimics the 20E dynamics previously determined by LC-MS (compare Sup. Fig. 2B and Sup. Vid. 1 with data from ^52^). These data show that ERE-NLS4xNG is a reasonable reporter of EcR, although the absence of NeonGreen fluorescence at the onset of the 3^rd^ instar (Fig. 2B and C) suggests that it has limited sensitivity (a more sensitive reporter based on EcR de-repression will be described below). Nevertheless, the pattern of fluorescence from ERE-NLS4xNG confirms the expectation that imaginal disc cells are exposed to an increasing level of 20E during the 3^rd^ instar and that proliferation arrest correlates with particularly high signalling activity. It is therefore plausible that different levels of 20E could trigger distinct effects on proliferation (promotion at low level, inhibition at high level).

**Figure 2:**
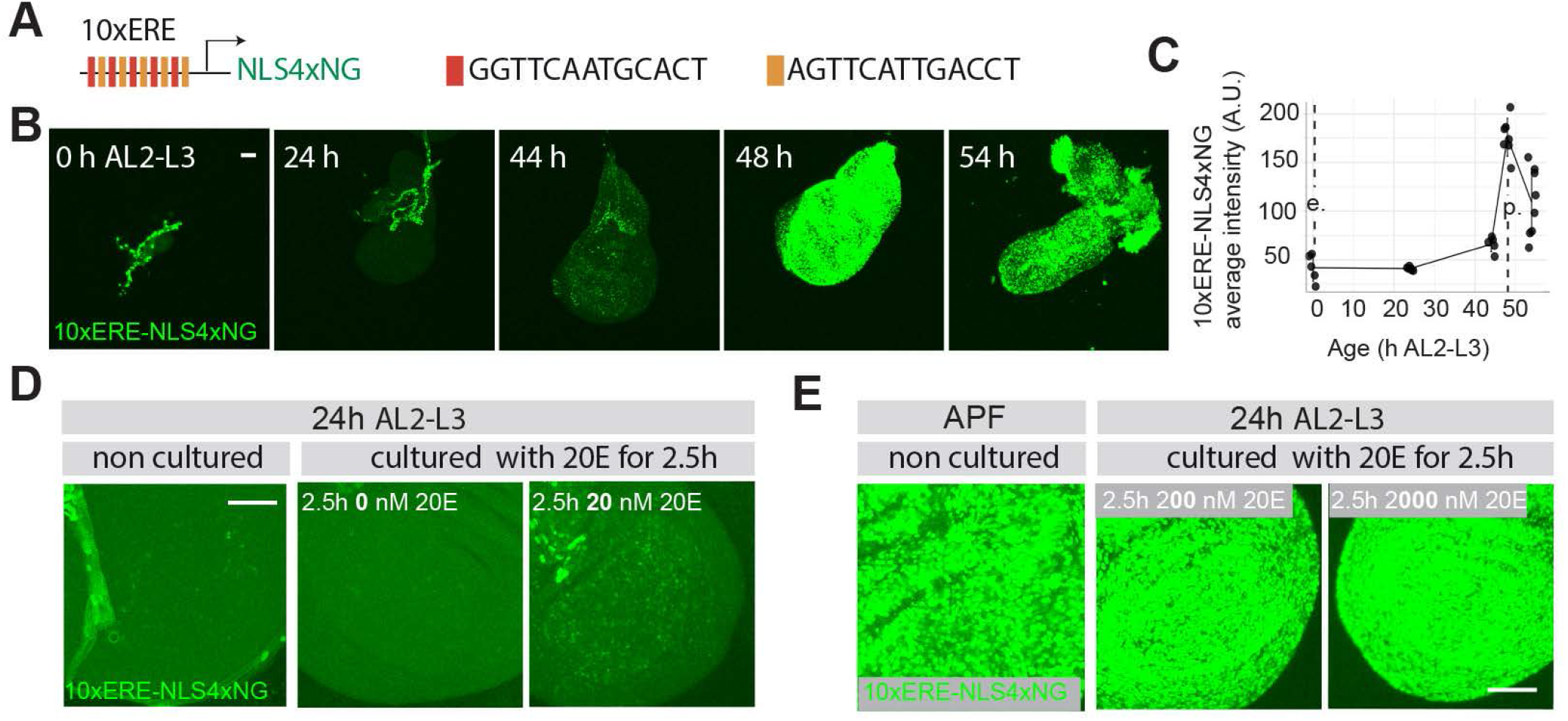
Effective 20E levels rise during the 3^rd^ instar. **A**. Schematic representation of the 10xERE-NLS4xNG reporter. **B-C**. Representative images and quantification of reporter fluorescence during the 3^rd^ instar and around the time of pupariation (all images taken under identical conditions). **D-E**. Estimation of in vivo 20E level at 24h AL2-L3 (mid L3) and at the onset of puparation. Reporter fluorescence intensity in discs explanted at 24h AL2-L3 and cultured for 2.5h with 20nM, 200nM, 2000nM 20E was compared to that in discs freshly explanted at 24h AL2-L3 or at pupariation. Each panel (D and E) shows images that are directly comparable. Scale bars represent 50 μm.

### 20E level determines whether EcR promotes or suppresses proliferation *ex vivo*

To estimate the dose response to 20E on growth and proliferation, we turned to *ex vivo* explants, which can be exposed to known concentrations of 20E. First, we calibrated the effective concentration of 20E that discs are exposed to *in vivo* by measuring the effect of 20E on ERE-NLS4xNG activity in explanted mid 3^rd^ instar discs. Immediately after dissection (non-cultured in Fig. 2D), weak but detectable fluorescence was present. In the absence of added 20E, this signal decayed to background level within 2.5 hour of culture (Fig. 2D middle panel), indicating that reporter activity is not sustained in the absence of 20E. By contrast, addition of 20nM 20E led to an increase in reporter activity (Fig. 2D right panel), which peaked at 2.5 hrs before dropping down (Sup. Fig. 2C and D). These observations suggest that, at the time of dissection (24h AL2-L3), *in vivo* concentration of 20E is lower than 20nM. The delay before peak expression could be caused by the time needed for transcription, translation and folding of NeonGreen, while subsequent decay of the signal could reflect suboptimal culture conditions and/or 20E depletion over time. Based on these results, we opted to measure the reporter’s dose response after 2.5 hrs in culture. Culture with 200nM or 2000nM of 20E (Fig. 2E middle and right panel) led to the same, strong signal, suggesting saturation of reporter activity above 200nM. Similarly strong reporter activity was seen in freshly explanted pupariating discs (Fig. 2E left panel), suggesting that, at this stage, *in vivo* 20E concentration is 200nM or higher. We conclude therefore that, *in vivo*, imaginal discs experience 20E at a concentration ranging from about 1-10nM at the onset of the growth period, to 200-2000nM at the time of pupariation. This agrees broadly with LCMS measurements, which suggest that peak 20E concentration at pupariation, is about 140 times higher than at the mid 3^rd^ instar ^52^.

Having established the range of 20E concentrations that imaginal discs are exposed to, we proceeded to assess how 20E affects proliferation in explanted mid 3^rd^ instar imaginal discs. As mentioned above, in the absence of 20E, proliferation ceased within 2.5 hrs in culture (Fig. 3A and Sup. Vid. 2). This could be rescued, in a dose-dependent manner, by addition of 10nM, 20nM or 40nM 20E in the culture medium (Fig.3A and Sup. Vid. 3-5). Proliferation was also sustained in EcR null imaginal discs without added 20E (Fig. 3B and Sup. Vid. 7-9), in accordance with the finding that EcR is dispensable for growth *in vivo*. The rate of proliferation in explanted EcR mutant discs was similar to that in explanted wild type discs treated with 20nM 20E (Fig. 3B), suggesting that sub-20nM 20E suffice to overcome the repressive influence of EcR on proliferation, while higher concentrations (20 and 40nM) could provide a further boost. However, upon exposure to 2000nM 20E, mitotic figures were no longer detected and early signs of eversion could be seen, despite the early stage (mid 3^rd^ instar) (Sup. Vid. 6). This effect of high concentration 20E is dependent on the EcR since no sign of eversion could be seen in mid 3^rd^ instar *EcR* mutant discs treated with 2000nM 20E (Sup. Fig. 3). Therefore, *ex vivo* as *in vivo*, 20E has a bimodal effect, promoting proliferation at low concentration, while preventing proliferation at high concentration. This suggest that 20E could trigger qualitatively distinct transcriptional responses at different concentrations.

**Figure 3:**
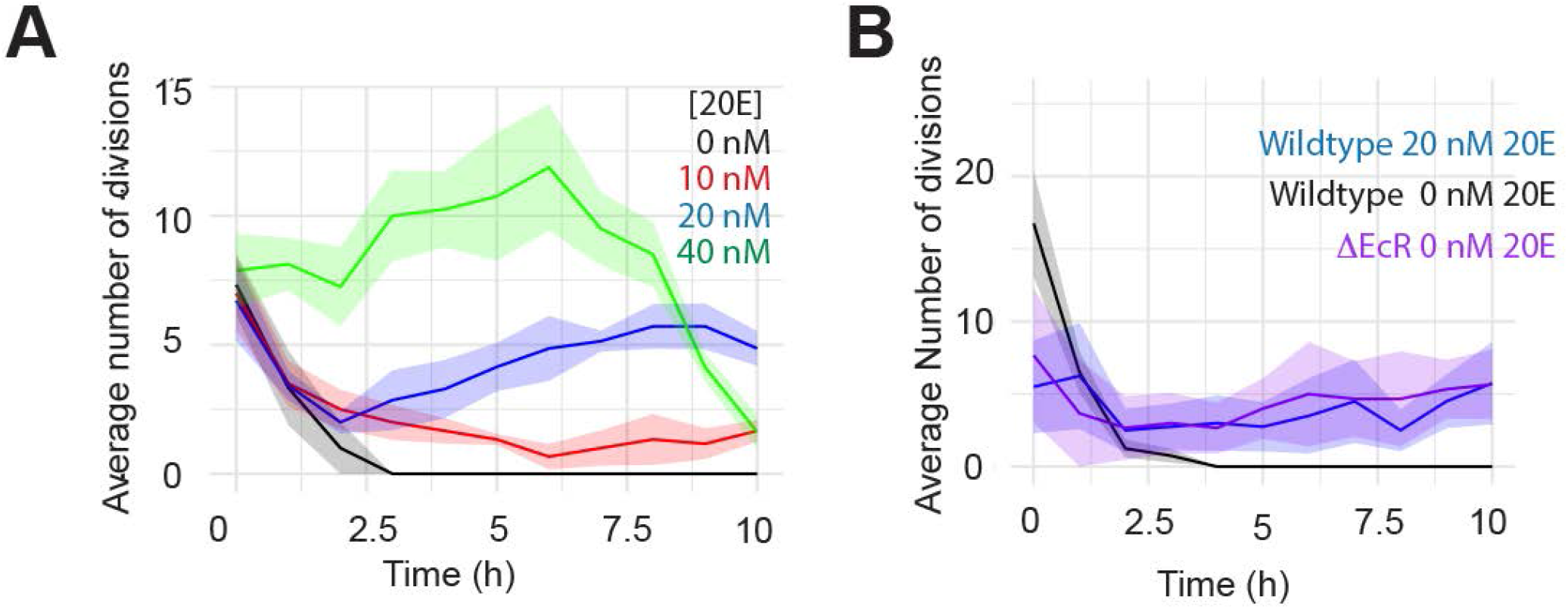
20E concentrations ranging from 10 to 40 nM promote proliferation ex vivo. **A**. Number of division measured in a region of interest (ROI) of wing disc explants cultured with the indicated 20E concentration. In the absence of 20E the discs stop proliferation (n=3) within 2.5h of culture. Adding 10 (n=6), 20 (n=7) or 40nM (n=8) 20E rescues proliferation in a concentration dependent manner. **B**. Inactivation of the EcR (n=3) allows explanted discs to proliferate at the same rate as wild type discs exposed to 20nM 20E (n=4), significantly faster than discs cultured in 0nM 20E (n=4). The average number of mitoses was calculated using a rolling 1 h period.

### Dose-dependent effects of 20E on transcriptional activity

To assess the transcription response to different concentrations of 20E, mid 3^rd^ instar wing imaginal discs were cultured for 2.5 hrs in 0, 20, 200 or 2000 nM 20E, and then processed for mRNA sequencing (Fig. 4A) and data analysis. As shown in Fig. 4B, a single principal component, which accounted for 89% of the variance, could reliably distinguish the four samples. As further evidence for the quality of the data, known targets of EcR signalling were found to be expressed in a concentration-dependent manner (Sup. Fig. 4A). This was confirmed by immunofluorescence for GFP-Br ^35,36^ and GFP-Blimp1^54^ (Sup. Fig. 4B-E). These observations give confidence that the transcriptional response of explanted mid 3^rd^ instar discs is physiologically relevant and warrants further analysis.

**Figure 4:**
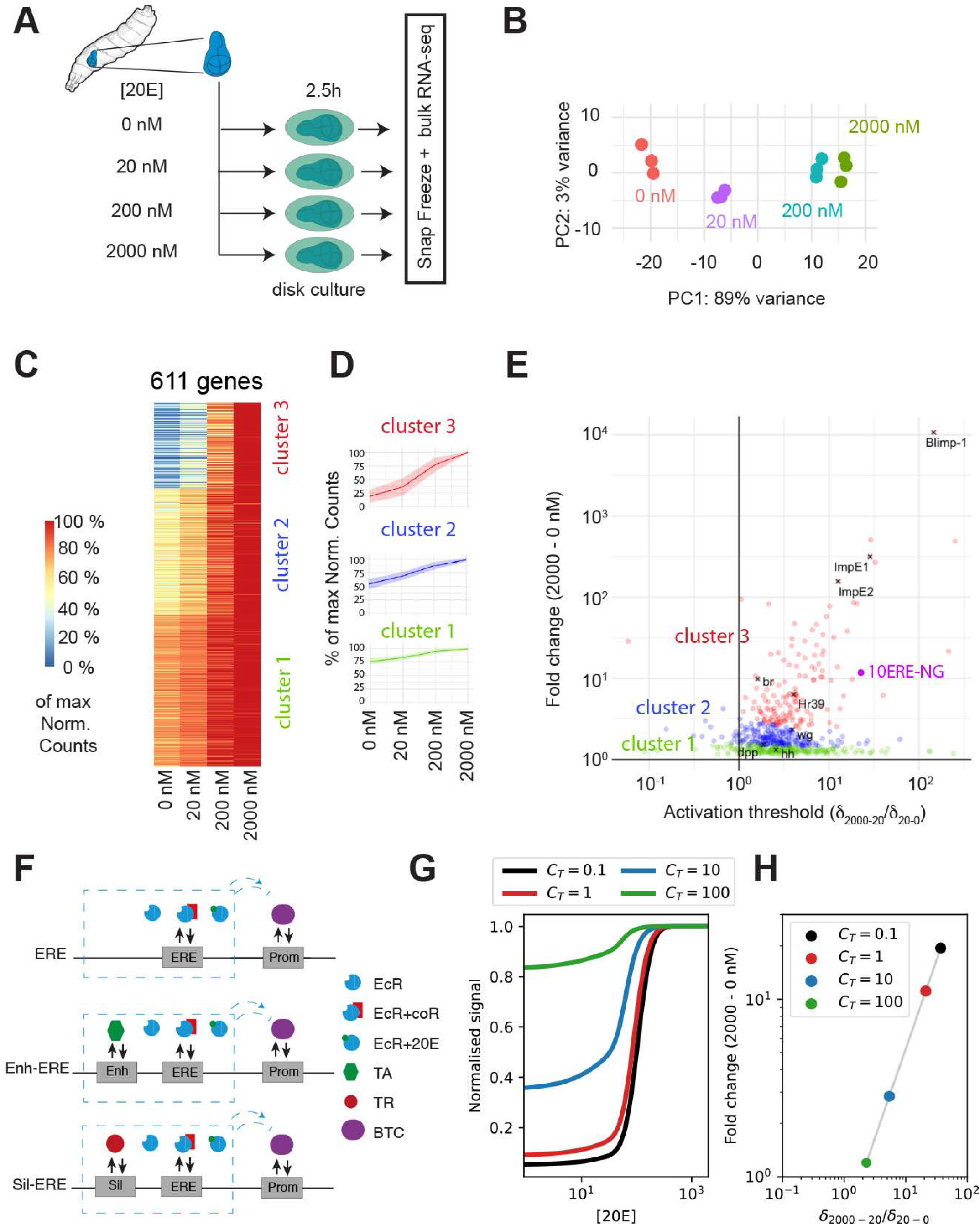
20E targets genes have various thresholds of activation. **A**. Experimental protocol to assess the transcriptional response to different 20E concentrations.**B**. Principal component analysis of the RNA-seq results shows clustering of the biological replicates. **C**. Transcriptional response of the 611 genes that are upregulated in response to increasing concentrations of 20E. Expression level is normalised to the highest value according to lookup table on the left. The gene responses were organised in three clusters as shown. **D**. Average response of each cluster, with standard deviation represented by a lightly coloured ribbon. **E**. Map displaying the extent of up-regulation of the 611 genes across the concentration range. The abscissa shows up-regulation in the 20-2000nM range relative to that in the 0-20nM range (a high value reflects a gene that is mostly activated at high concentration). This is plotted in relation to the overall fold change of expression. Genes are colour-coded according to the cluster they belong to. **F**. Diagrammatic representation of regulation of three hypothetical target genes considered by the thermodynamic model, which differ by the presence or absence of basal enhancer (Enh) and silencers (Sil), to which transcriptional activators (TA) or transcriptional repressors (TR) can bind. In the model, EcRs bound to ERE act as activator of BTC binding to the promoter when bound to ecdysone, and as repressor when bound to a corepressor. **G**. Transcriptional activity as a function of ecdysone concentration predicted by the thermodynamic model, normalised to the maximum, for a gene regulated by an ERE (red curve), in the presence of a basal enhancer of increasing strength (blue and green curves) and in the presence of a basal silencer (black curve). The presence of the enhancer results in an increased finite baseline activity at low 20E, as well as a decrease of the threshold ecdysone concentration at which the genes expression level changes. *κ*_*P*_ = 0.1, 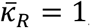, *C*_*ER*_ = 0.1, *C*_*EA*_ = 10. **H**. Predicted relationship between fold change increase in transcriptional rate (y axis) and relative increase in gene expression at 2000nM-20nM vs 20nM-0nM ecdysone concentration (x axis), as the strength of a basal enhancer is varied. Colored dots correspond to curves in panel G.

As a first level of analysis, we used DESEQ2 to identify genes whose changes in expression could be explained by dose dependence on 20E (see Material and Methods). Among the 1489 resulting genes, we retained only those changing monotonically and with a total read number exceeding a small arbitrary threshold (see Methods for further details). This first selection identified 611 genes that are upregulated in an 20E concentration dependent manner and 635 genes that are downregulated. Since 20E-bound EcR acts as a transcriptional activator ^12^, we surmised that the downregulated genes are repressed indirectly ^53^. Indeed, these genes had relatively few EcR binding sites in the vicinity of their transcription start site (Sup. Fig. 4F-G). In contrast, all upregulated genes were characterised by an enrichment of canonical EcR binding sites (Sup. Fig. 4F-G). We therefore chose to restrict subsequent analysis to the 611 upregulated genes, which are most likely controlled directly by EcR and 20E.

For each gene, reads were normalised to the highest value at any of the four concentrations, thus allowing the different dose responses to be compared despite wide ranges in expression levels. The results were then plotted on a heatmap (Fig. 4C). K-medoids clustering of normalised gene expression revealed that 20E target genes could be classified in three clusters according to the shape of their response (Fig. 4D). Thus, cluster 1 genes are expressed in a dose-dependent manner at all 20E levels, but with significant basal expression in the absence of 20E. Cluster 3 genes (red), by contrast, do not respond to low concentration 20E and display concentration-dependent behaviour only at high concentrations. These will be referred to as high threshold genes by analogy to the genes that respond only to high level morphogen signalling. Finally, cluster 2 (blue) showed an intermediate response. The behaviour of representative genes from each of the three clusters is shown in Sup. Fig. 4H. To determine, for any given gene, whether the dynamic range of the response lies mostly at high or low 20E concentrations, we devised the δ_(2000−20)_/δ_(20−0)_ ratio, which compares the fold change in gene expression between 0 and 20nm to that between 20 and 2000nM. This parameter was plotted relative to the overall fold change (between 0 and 2000nM) (Fig. 4E). In the resulting response map, high threshold cluster 3 genes (red dots) appear mostly on the upper right side. By contrast, cluster 1 genes (green) tend to display a relatively low overall fold change and therefore appear at the bottom of the map. This map shows a weak but significant correlation (R=0.26) between δ_(2000−20)_/δ_(20−0)_ and the overall fold change (δ_(2000−0)_). By comparison no such correlation could be seen with a dataset of randomly generated virtual genes (R= -0.054) (Sup. Fig. 4I and J), suggesting that the correlation between overall fold change in gene expression and the tendency to respond only to high 20E concentrations, as seen in the feature map, is genuine. In conclusion, RNAseq analysis reveals a range of transcriptional responses, which could underly the bimodal effect of 20E on proliferation. Thus, we expect genes involved in termination of proliferation to fall in the upper-right side of the response map (mostly expressed at high 20E concentrations). Pro-proliferation genes could be possibly found in rest of the map, i.e. in the lower-left side (active at low 20E concentrations), although we cannot exclude the possibility that targets not considered by our analysis could mediate the pro-proliferation activity of 20E (see discussion).

### Emulating EcR target gene behaviours *in silico* and *in vivo*

The 10xERE-NLS4xNG reporter, which receives its inputs only from EcR and 20E is unlikely to recapitulate the range of dose responses of EcR target genes. Calculation of the δ_(2000−20)_/δ_(20−0)_ ratio places this minimal reporter within cluster 3 (high-threshold targets, Fig. 4E purple dot). What are the regulatory features that would allow a reporter gene to mimic the range of behaviour seen with natural EcR responsive genes? Cluster 1 genes are characterized by non-zero baseline activity. They must therefore receive a positive input from a separate enhancer element. At the other extreme, high threshold genes remain silent in the 0-20nM range of 20E concentrations. This could be achieved by a silencer element that only allows expression at high concentrations of 20E.

To mathematically explore how such simple elements would alter the response of 10xERE-NLS4xNG, we devised a coarse-grained thermodynamic model of transcriptional regulation that combines the effect EREs to those of additional basal enhancer/silencer elements (see Mathematical Modeling in Material and Methods). In this model, an average transcriptional activity is derived from the probability of binding of the basal transcriptional complex (BTC) to the promoter of the gene ^55-60^. The affinity of the BTC for the promoter depends on whether the EcR is associated with its co-activator or co-repressor, as well as on the presence of an additional basal activator or repressor. Competitive binding of the co-activator or co-repressor to EcR was incorporated in a simple model that tracks the probability of the possible complexes (see schematic of Fig. 4F and Sup. Fig. 5A, B) ^11-19^. This model successfully recapitulates the previously reported “sponge effect” of a diffusible EcR fragment lacking its DNA binding domain ^61^ (Sup. Fig. 5C) and predicts first order Hill-type functional forms for *P*_*act*_(*E*) and *P*_*rep*_(*E*), the probabilities that the free EcR is in its activating and repressing form, respectively, as a function of the concentration of 20E (denoted as *E*). Next, we derived an expression for the normalized transcriptional activity *A* of a gene regulated by a single EcR and additional basal transcriptional activator or repressor:

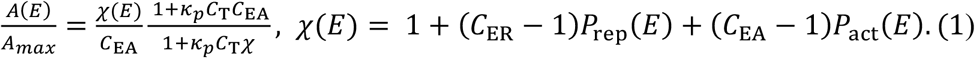

Here, the cooperativity coefficients (*C*_ER_, *C*_EA_, *C*_T_), modulate the basal affinity of the BTC for the promoter *κ*_*p*_, taking into account the configuration of EcR, i.e. in its repressive form (*C*_ER_< 1), activating form (*C*_EA_> 1) or neutral form, and/or the effects of an additional basal transcriptional activator (*C*_T_> 1) or repressor (*C*_T_< 1). The function *A*(*E*) in equation 1 is characterised by a finite baseline value that increases with *C*_T_ (Fig. 4G). By adjusting *C*_T_ (the strength of the basal transcriptional activator/repressor), the model can reproduce the characteristic normalised response of cluster 1, 2, and 3 genes (Fig. 4G). Increasing *C*_T_ indeed reduces the overall fold change of activation by 20E, defined as the ratio of expression levels at high and low E20 concentration (Fig. 4G). Across the values of *C*_T_, the predicted overall fold activation correlated positively with the predicted δ_(2000−20)_/δ_(20−0)_ ratio (Fig. 4H), as observed with real 20E target genes (Fig. 4E). The thermodynamic model explains this correlation through the dual effect of a basal enhancer. The enhancer reduces the overall fold change of activation and also increases the effective affinity of 20E for DNA-bound EcR (as opposed to free EcR). This is because the enhancer stabilises the BTC at the promoter, which in turn thermodynamically favours 20E binding to DNA-bound EcR in the same way that 20E-EcR favours BTC recruitment to the promoter (Mathematical Modeling in Material and Methods). Conversely, silencers are predicted to increase the overall fold change of gene expression and increase the threshold of activation by 20E to higher 20E concentration (Fig. 4H). This predicted dual effect of basal enhancers and silencers provides a simple explanation for the correlation between the fold change increase of gene expression in response to 20E and their threshold of activation (Fig. 4E). Overall, our mathematical analysis demonstrates that combining EREs with a constitutive enhancer or silencer is needed to recapitulate the complete range of responses of EcR target genes.

We next proceeded to build synthetic reporters based on the principles outlined above. To mimic high threshold target genes, we inserted upstream of 10xERE two copies of a silencer element from the *brinker* gene, Brk^S 62^, which has been shown to mediate constitutive repression in the prospective wing (Schematic in Fig. 5A). Activity of the resulting 2xBrk^S^-10xERE-NLS4xNG reporter was first assessed in transgenic imaginal discs at different developmental stages (Sup. Fig. 6A). A fluorescence signal could only be detected at the end of the 3^rd^ instar, at the time of pupariation, suggesting that this reporter only responds to high level 20E. A control transgene comprising mutated EREs (EREs*) was silent at all stages (Sup. Fig. 6B), confirming that this reporter’s response to high 20E levels depends on functional EREs. We next assessed the activity of 2xBrk^S^-10xERE-NLS4xNG in cultured mid 3^rd^ instar imaginal discs treated with 20E at different concentrations. A fluorescence signal was only detected after treatment with high 20E concentrations (200 nM or more), while the control transgene had no activity, even at high 20E concentration (Fig. 5A and B). We conclude that the high threshold behaviour of the minimal ERE reporter can be exaggerated (pushed to the upper-right side of cluster 3 genes in the response map) by addition of a constitutive silencer.

**Figure 5:**
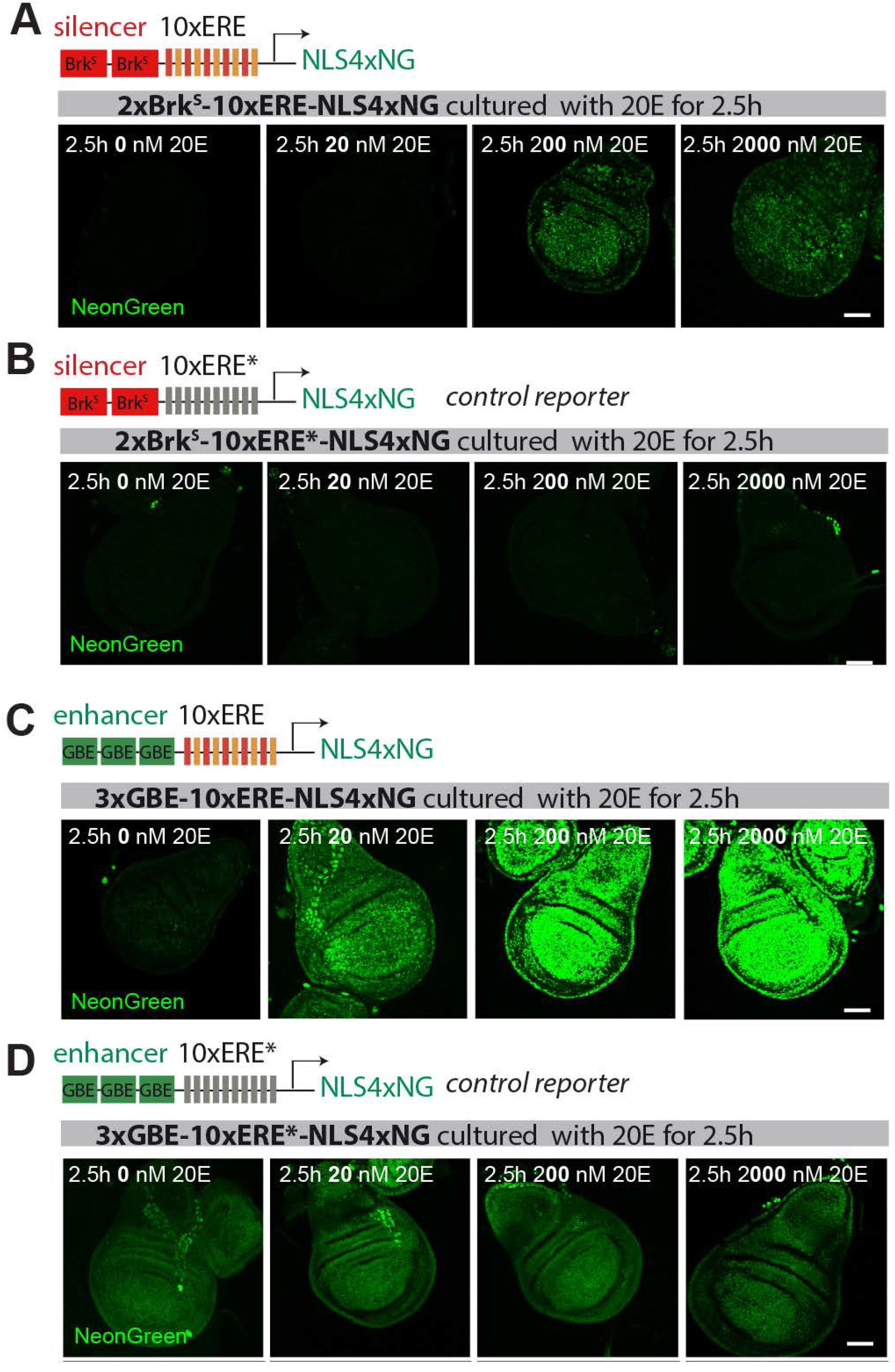
Emulation of various responses to 20E in synthetic reporters. **A-B**. Addition of a silencer (2xBrk^S^) raises the concentration of 20E needed to trigger activation by ERE. While 20nM 20E suffice for detectable activation of 10xERE-NLS4xNG (see Figure 2B), 200nM are needed to activate 2xBrk^S^-10xERE-NLS4xNG. Therefore, this reporter emulates a high threshold target gene. A control reporter with mutated ERE is not activated at any concentration (B). **C-D**. Addition of an enhancer (3xGRE) to 10xERE-NLS4xNG raises baseline activity, leading to weak, albeit detectable signal, even in the absence of 20E. Note that in the absence of 20E, the reporter is less active than the control reporter (mutated ERE) because of repression by unliganded EcR. The 3xGBE-10xERE-NLS4xNG reporter emulates the expression expected from pro-proliferative genes (responding to all physiological concentrations of 20E).

To mimic genes located at the other end of the map (expressed across the range of 20E concentrations), we used Grainy head binding elements (3xGRE), which confer baseline enhancer activity in the prospective wing ^63^, in combination with EREs. In cultured mid 3^rd^ instar discs, the resulting reporter was expressed in a 20E concentration-dependent manner but with non-zero activity at 0nM 20E (Fig. 5C), as seen with cluster 1 genes. The corresponding control transgene (with ERE*) was expressed at an intermediate constitutive level at all concentration of 20E (Fig. 5D), as expected. At 0nM 20E, this constitutive signal was higher than that of 3xGBE-10xERE-NLS4xNG (with wildtype EREs), indicating that, in the absence of 20E, EcR suppresses the activity of the baseline enhancer. At 20nM and above, the situation is reversed with the GBE-ERE combination overtaking the baseline enhancer (GBE only) (Fig. 5D and Mathematical Modeling in Material and Methods). Similar conclusions can be drawn from reporter activity observed at increasing times after the L2-L3 transition (Sup. Fig. 6D). Expression of this reporter at 24h AL2-L3, but not at 44h AL2-L3, is below the baseline activity from the constitutive enhancer, highlighting once again the contribution of 20E in relieving the default growth-repressing activity of EcR during the growth phase of imaginal discs. Overall, the above results indicate that 3xGBE-10xERE-NLS4xNG emulates relatively well the behaviour of low level 20E target genes.

We then tested our thermodynamic model by comparing its prediction to the experimental responses of the three transgenes, through a numerical fit that allowed us to extract its free parameters (Fig. 6D). The results showed that all response curves could be explained by the thermodynamic model. The best fit parameters revealed that the enhancer-bound basal activator has a weaker effect on transcription (*C*_TA_ ≃ 2) that 20E-bound EcR (*C*_EA_ ≃ 13.2); indicating that the EcR strongly activates transcription at high 20E concentration (see details in the Mathematical Modeling in Material and Methods). In combination with the low basal affinity of the BTC for the promoter (*κ*_*P*_ ≪ 1), this leads to a large overall fold change in response to 20E. Model fitting also implies that EcR acts as strong repressor at low ecdysone concentration (*C*_ER_ ≃ 0.09) and that the repression is lifted at an ecdysone concentration of ∼ 5nM, broadly consistent with the observation that low ecdysone concentrations of 10, 20nM can restore proliferation (Fig. 3A, B). We found that the transgene response curves were best explained by models involving cooperativity between EcR bound to several ERE sites, albeit at the cost of additional assumptions needed to take cooperativity into account (Fig. S5D-F and Mathematical Modeling in Material and Methods). Overall, our *in vivo* and *in silico* results show that simple rules can reproduce the behaviour of a wide range of EcR target genes, including that of synthetic reporters.

**Figure 6:**
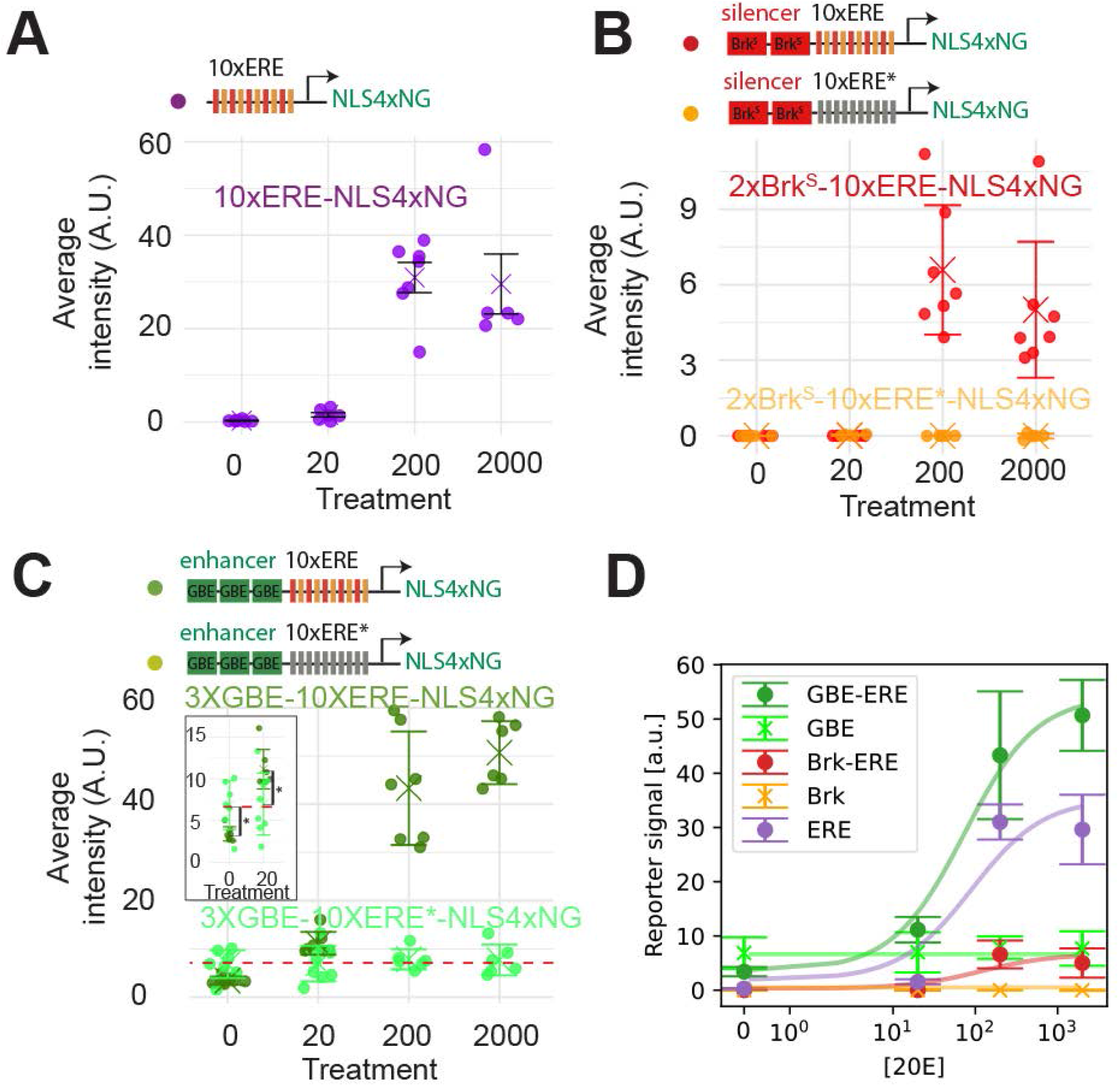
The thermodynamic model predicts cooperativity between EcR and the basal activator. **A-C**. Quantification of reporter activity (NeonGreen fluorescence) in transgenic wing disc cultured for 2.5 h for different concentrations of 20E, expressed in nM. Error bars represent standard deviation. T-tests were performed in C. * p<0.05. **D**. Experimental data from A-C and fitted curves from the thermodynamic model for different constructs, assuming for simplicity that the activation probability of the set of EREs as a function of ecdysone equals that of a single ERE (more detailed descriptions accounting for interactions among EREs are explored in Sup. Fig. 5F). Ecdysone levels are plotted on a symmetric logarithmic scale. Parameters: *κ*_*P*_ ≃ 0.03, 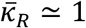, *C*_ER_ ≃ 0.1, *C*_EA_ ≃ 13, *C*_T2_ ≃ 0.1, *C*_TA_ ≃ 2 *k*_*T*_ ≃ 131 a. u. (see Mathematical Modeling in Material and Methods).

## DISCUSSION

In this paper, we reconcile two seemingly opposite views on the effect of systemic ecdysone in the control of tissue proliferation. We show that, during the third larval instar of Drosophila, the main growth period of wing precursors, ecdysone is required for proliferation and growth, while at the end of larval life, the pulse of ecdysone that triggers pupariation causes proliferation arrest. These opposite effects can be accounted for by a differential response to the different levels of ecdysone that are present at these two stages. Thus, in cultured mid 3^rd^ instar wing imaginal discs, addition of 10-40nM 20E sustains proliferation, while higher concentrations of 20E (e.g. 200nM) lead to rapid proliferation arrest. Remarkably, the ecdysone receptor is not strictly required for the pro-proliferative effect of ecdysone; proliferation is sustained both in vivo and ex vivo in EcR null imaginal discs, even in the absence of ecdysone. We therefore suggest that during the growth phase of imaginal discs, the EcR acts as a brake to proliferation via its unliganded repressor activity, and that this brake is progressively released and overcome by the rising level of ecdysone. At much higher concentrations such as those that are present at pupariation, ecdysone does not merely derepress the EcR but, in addition, activates a gene expression programme that triggers morphogenesis and proliferation arrest. The basal signalling activity seen in the absence of EcR and its ligand is reminiscent of other signalling pathways where removal of the main transcriptional mediator leads to weak though significant ligand-independent signalling activity (e.g. TCF for Wnt signaling or Gli for Hedgehog signaling) ^59,64-67^. This arrangement achieves a greater dynamic range by ensuring lowering signaling activity in the absence of ligand. In the case of EcR, it appears that three functional outputs are generated: (1) the absence of ligand prevents proliferation; (2) low ligand levels mimic the default pro-proliferative state (same as in the absence of EcR); (3) additional signalling due to higher ligand levels terminates proliferation. Thus, depending on its systemic concentration, the same hormone has opposite effects on proliferation.

To further explore the dose-dependent activity of ecdysone, we turned to RNAseq analysis of explanted imaginal discs cultured exposed to a range of concentrations. Many genes were found to be activated in a dose-dependent manner, but they differed in the shape of their response curve. Some genes were primarily expressed at high 20E concentrations (cluster 3), while others were expressed across the entire range of physiological concentrations (clusters 1 and 2). Among the former, we found many genes previously shown to be activated at pupariation, when ecdysone levels are relatively high, e.g. Blimp1, ImpE1, ImpE2 and ImpL2 ^68^, validating our analysis. Genes involved in proliferation arrest are likely to be found among these high threshold genes. However, so far, we have not been able to identify a gene that, on its own, is sufficient to prevent proliferation upon overexpression at mid 3^rd^ instar, presumably because proliferation arrest requires the coordinated activation of multiple high threshold genes. At the other end of the spectrum, among genes expressed at low ecdysone concentrations, were *wg, dpp, hh* and *Egfr*, which are required for imaginal tissue growth. These and also genes involved in Hippo and mTor signaling, have been shown previously to be regulated by ecdysone ^28-31^ either directly, as suggested by EcR ChIP-seq analysis ^48^ or indirectly via modulation of Taiman (Tai), an EcR cofactor ^69,70^. We note however that these genes are expressed in the absence of 20E, when proliferation is not sustained. Perhaps, they only stimulate proliferation in concert and over a combined threshold. Alternatively, additional 20E responsive genes not uncovered by our analysis (perhaps because they are not directly activated by 20E) may be required. A molecular understanding of proliferation control by 20E remains a formidable challenge. It is worth pointing out that no EcR target genes were found to be expressed only in pro-growth concentration range. We suggest therefore that the anti-proliferative genes that are expressed at high concentration must override the effect of pro-growth target genes (see diagram Fig.7).

**Figure 7:**
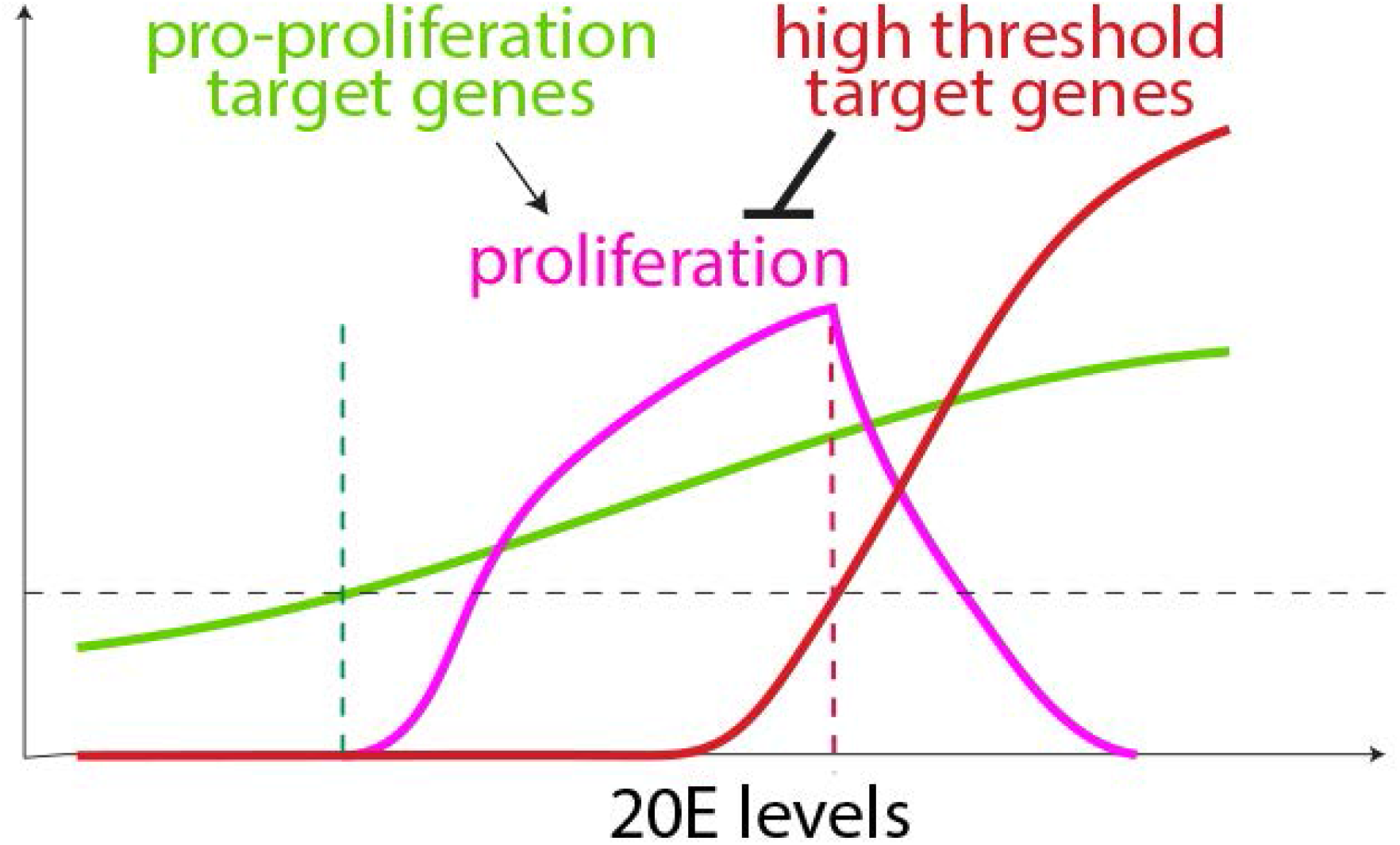
Model: Control of cell proliferation by different levels of 20E. **A**. Pro-proliferative genes increase in response to all levels of 20E, while high threshold targets genes (anti-proliferative) are only activated at high 20E concentrations. High threshold anti-proliferative genes are proposed to dominantly suppress the activity of pro-proliferative genes. We suggest that the presence of basal enhancers or silencer modulates the activity of EREs to determine the dose-response of the two classes of genes, as described in the text. The repressive function of unliganded EcR would guarantee the inhibition of target genes in the absence of 20E, perhaps explaining the requirement of 20E for growth.

To understand how a simple regulatory element can achieve qualitatively distinct dose responses, we took a synthetic approach, first in silico and then in vivo. Our results suggest that the 20E threshold required for the activation of target genes is affected by the architecture of their cis-regulatory region, the presence or absence of basal enhancers or silencers in particular. This combination of regulatory elements can lead to modulation of the response of gene expression to ecdysone concentration. Since ecdysone concentration increases over time during development, these responses can in principle give rise to various patterns of temporal activation of genes, analogous to spatial morphogen gradients triggering spatial domains of target gene expression (Sup. Fig. 5E ^61^). However, other features are also likely to be relevant. For example, McKay and colleagues have shown that chromatin accessibility to EREs is a key determinant of the EcR’s response ^71^. Moreover, several targets of ecdysone signalling are known to modulate EcR activity (for example Eip78C^72^), highlighting the importance of feedback control. These features could be incorporated in an expanded model that make the activity of the basal enhancer, or of liganded EcR dependent on current or historical 20E levels. Nevertheless, in its current form, our model shows that a small number of regulatory elements can account for the spectrum of responses to 20E.

In summary, our work explains how a given type II nuclear receptor can drive opposite cellular responses depending on the levels of its ligand. We showed that the presence of a basal enhancer or silencer could determine whether a target gene responds only to high hormone concentrations or to a broader range that also includes low concentrations. We further suggest that a bimodal response can be achieved if the high threshold genes suppress the activity of the low threshold genes. The bimodal nature of EcR signalling could be relevant to vertebrate type-II nuclear receptors such as the retinoic acid receptor ^73,74^or the thyroid receptor ^75,76^. Moreover, our work highlights the possibility that inactivation of the receptor, e.g. with a chemical degrader, may not achieve the same objective as hormone depletion.

## Supporting information

Mathematical Modelling

## ACKNOWLEDGMENTS

We are grateful to Nic Tapon for allowing us to use 4xNeonGreen in our reporter constructs, Natalie Dye for hosting GPM and teaching him how to image wing imaginal discs for extended periods and Yifan Zhao for help with some experiments. We also acknowledge Giorgos Pyrowolakis for help with the design of synthetic reporters and Nic Tapon for comments on the manuscript. The Crick’s Advanced Light Microscopy Facility (CALM) provided help with imaging and the Advanced Sequencing Facility performed library preparation and mRNA-seq.

## AUTHORS CONTRIBUTIONS

This project was conceived by GPM and JPV. GPM constructed the ECR^KO^, the different synthetic reporters, performed the fly work and imaging, and carried out the bioinformatic analysis of the bulk RNA-sequencing. CA built the EcR^CKO^ fly line. BA devised the 4xNeonGreen-encoding plasmid used for the reporters. LC and GS devised the mathematical model, LC performed calculations and contributed to experimental design with advice from GS. GPM wrote the first draft except the parts related to the mathematical modelling, which were written by LC. The draft was subsequently modified and edited by all the authors.

## MATERIAL AND METHODS

### Fly stocks and husbandry

Flies were reared in standard cornmeal/agar media at 25C in a Sanyo incubator with 12h light/dark cycles.

### Developmental curves

Flies were allowed to lay eggs for 4 h intervals between 8:00 and 20:00. 2 × 30 L1s for each of these three plates were then transferred to fly vials (six tubes in total) and pupae formation was scored every 4 hours between 8:00 and 20:00.

### Fly Genotypes

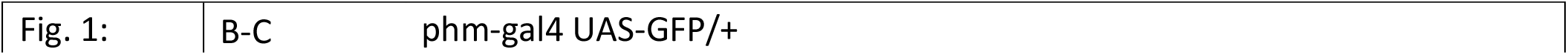
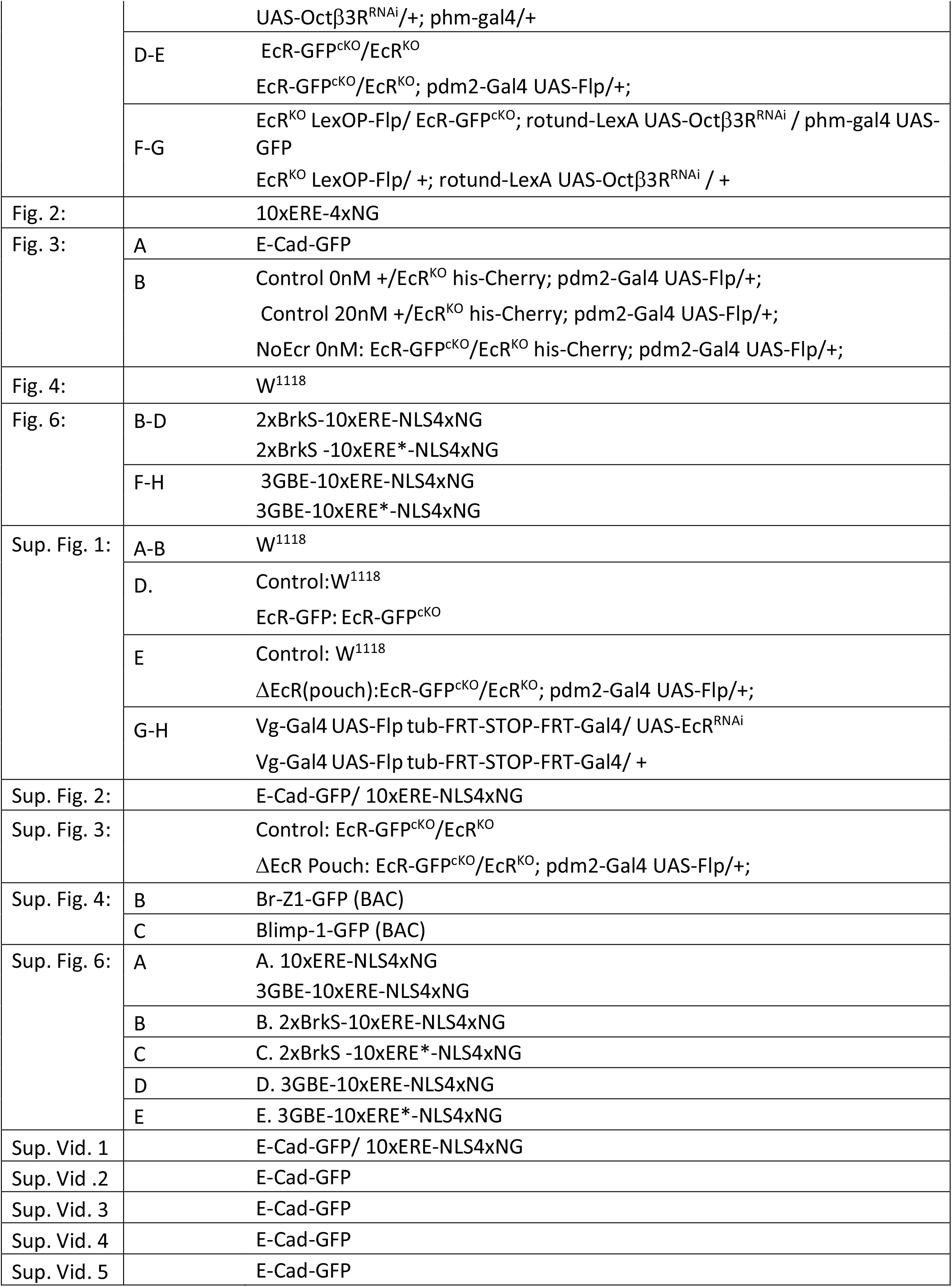
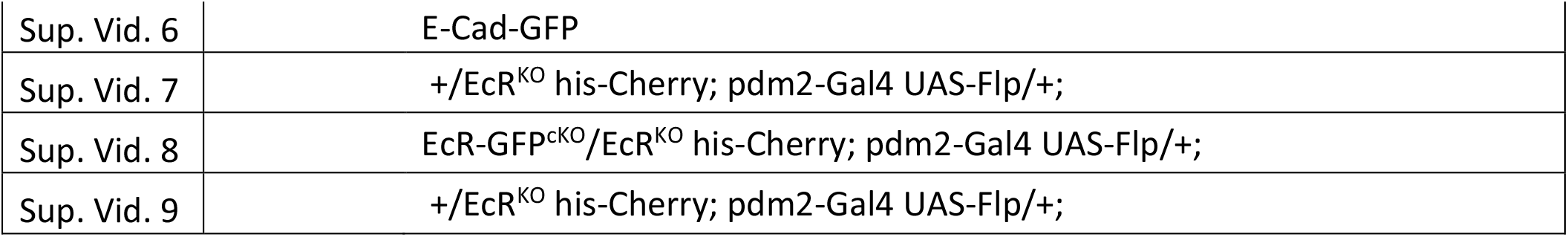

### Wing disc culture and Imaging

Dye medium was prepared as described in ^30,77^. Briefly, Grace’s medium (Sigma, G9771) containing 5mM BisTris had its pH adjusted to 6.6-6.7. Prior to each experiment, it was supplemented with 5% FBS (ThermoFisher/Invitrogen, 10270098), 1x Pen/Strep (Sigma P4333, 100x stock solution) and different concentrations of 20E (Sigma, H5142).

Live imaging experiments were performed by mounting wing discs in an uncoated ibidi 35mm imaging dish (Ibidi, 81141) as described in ^38^. We used a Nikon CSU-W1 Spinning Disk, to image the disc every 5 minutes with a Z-interval of 0.75 μm and using the 60x objective.

After performing a max projection, the E-Cad or Histone signal was used to manually track mitosis in a region of interested (ROI). This ROI was determined such as its size was comparable between different conditions and replicates.

For ex vivo culture, 24h AL2-L3 discs were dissected and incubated for 2.5h at 25 °C in Dye medium supplemented with different 20E levels. They were then fixed in 4% formaldehyde (Pierce 28906) for 45 minutes.

For NeonGreen fluorescence, Br-Z1-GFP and Blimp-1-GFP immunofluorescence quantifications a custom code was used to calculate the intensity of the signal inside the nucleus (marked with DAPI) minus the noise measured outside of the nucleus.

Data presented in Sup. Fig. 6 was used to normalise data from the 10xERE-NLS4xNG and the 3xGBE-10xERE-NLS4xNG/3xGBE-10xERE*-NLS4xNG/2Brk^S^-10xERE-NLS4xNG/2Brk^S^-

10xERE*-NLS4xNG reporters, which were acquired under different conditions.

Expression of Br-Z1 and Blimp-1 proteins was inferred from staining knock-in strains with chicken anti-GFP (Abcam Ab13970 1:500) and an anti-chicken secondary antibody (Invitrogen A-21437 1:1000).

### Pupal imaging

A single focal plane was recorded every 20 min on a live cell imaging Nikon LTTL 1 (4x objective). For quantification, an ROI was manually selected as shown in the Sup. Videos 2-9, and the average fluorescence intensity was measured inside this ROI at every time point.

### Volumetric Analysis

Volume quantifications were obtained from staged wing discs fixed for 45 minutes in 4% PFA (Pierce 28906), stained with Vectashield® (Vector labs, H-1200-10), mounted in agar as described in ^38^, and imaged with an upright Leica SP5 confocal microscope. Since Vectashield® DAPI non-specifically stained at low levels the whole wing imaginal disc tissue, it was possible to use Imaris to generate 3D reconstructions and measure tissue volume.

### Molecular Biology and Cloning

The *EcR*^*CKO*^ line was generated by removing the last 4 exons common to all the EcR isoforms and replacing them with an attP sequence. The two CRISPR target sites used were located 526bp upstream of the four last common exons, and 1 bp after the stop codon in the last exon. We then re-inserted a genomic fragment containing the last 4 exons, eGFP before the stop codon and 2 kb of 3’UTR.

*EcR*^*KO*^ was made by removing the last 3 exons using a CRISPR target site located 132 bp upstream of the last 3 exons, and another located 32 bp after the last exon.

For the various NeonGreen reporters, we used the GeneArt Gene Service from Thermo Fischer (https://www.thermofisher.com/uk/en/home/life-science/cloning/gene-synthesis/geneart-gene-synthesis.html) to synthetise either 3xGBE, 2xBrk^S^, 10xERE, or 10xERE*. The sequences were then inserted in the relevant order using restriction digest and ligation. All sequences and details are available upon request.

### Sample preparation for RNAseq

24h AL2-L3 wing discs were incubated for 2.5h at 25°C in Dye medium (see wing disc culture) supplemented with different concentrations of 20E (Sigma, H5142). Discs were then snap-frozen in dry ice. After all the samples were collected, RNA was extracted with an RNeasy Mini Kit (Qiagen, 74104). 1.2-2.1 μg were used as a template to generate a sequencing library with the NEBNext Ultra II Directional PolyA mRNA (NEB, E7760S). The Advanced Sequencing Facility of the Francis Crick institute used an Illumina HiSeq 4000 to perform single end 1 × 75 bp sequencing.

### RNA Sequencing Analysis

The reads were aligned to BDGP6, reference genome (ensemble release 84), using HISAT2(v. 2.1.0) ^78^. SAMtools (v. 1.13) ^79^ allowed to first transform the HISAT2-generated SAM files into BAM files and then to sort and index them. FeaturesCounts from the package Subread (v.1.6.4) ^80^ was used with the options -t exon \ -g gene_id \ --primary to count reads mapping with features. All these tasks were parallelised using SLURM ^81^. Principal component analysis was performed on the vst transformed (option blind=TRUE) raw data, and plotted using the plot PCA function of DESeq2^82^.

DESeq2 was used to determine the genes that displayed a change in expression that could be explained by changes in ecdysone levels (Likelihood ratio test p adjusted value<0.001). To simplify the analysis, we decided to ignore genes that had a total average count (from the 4 experimental conditions) below 500. This was based on the assumption that such low expression level was unlikely to have biological significance. We also ignored the genes that did not change monotonically between 0, 20 and 200 nM, as they were more likely to be indirect targets of EcR. The average counts (normalised using DESeq2’s median of the ratio methods) were further normalised to the maximum value for each gene, and then separated into clusters using k-medoids clustering from the R cluster package (v.2.1.4 ^83^). The optimal cluster number was determined using the elbow method on a graph representing the total intra-cluster variation in function of the number of clusters.

For the randomly generated data, we pooled the dataset containing all the genes affected by ecdysone (going up and down at all concentrations). We then calculated the mean and standard deviation across the whole data and used a gaussian distribution with the same mean and standard deviation to generate an artificial dataset composed of 50 000 synthetic genes. The same filtering used previously to select the upregulated genes was used to filter down this randomly generated dataset. This led to the 8371 genes presented in the Supplementary Figure 4 I.

Cytoscape (v. 3.9.1) ^84^ plugin Iregulon (v.1.3) ^85,86^was used to identify the transcription factors most likely to bind the regulatory region of genes positively and negatively regulated by ecdysone. The following parameters were used: motif collection, 10K; species and gene nomenclature, Drosophila melanogaster; Flybase names; region-based specific parameters, Overlap fraction 0.4, 10kb upstream, full transcript and 10kb downstream; recovery enrichment score threshold, 2.5; ROC threshold, 0.001; rank threshold 5000, TF prediction minimum identity, 0.0; FDR, 0.001.

## FIGURE LEGENDS

**Sup. Fig.1 related to Figure 1. Wing growth and genetic tools to assess role of EcR**.

**A**. Representative images of the volumetric reconstruction of L3 wing discs and pupal wings.

**B**. Volume quantification during growth. **C**. When 20E production is impaired (Δ20E; phm>OctΔ3^i^), larvae do not pupariate and continue growing. **D**. Schematic representation of the EcR locus showing the three isoforms A, B1 and B2. In EcR-GFP^cKO^. A GFP tag was inserted as shown and FRT sites were inserted to flank the last four exons, which are common to all isoforms. **E**. Data showing that EcR-GFP^cKO^, which is homozygous viable does not affect developmental timing. **F**. The product of EcR-GFP^cKO^, detected by GFP fluorescence, localizes to the nucleus. **G**. Inactivation of EcR-GFP^cKO^ (pdm2>FLP EcR-GFP^cKO^ / EcR^KO^) had no impact on developmental timing. **H-J**. Whole wing knockdown of EcR in the whole wing with constitutive Vg-Gal4 driving an RNAi transgene (Vg-Gal4 UAS-Flp Tub-FRT-STOP-FRT-Gal4/ UAS-EcR^RNAi^) leads to overgrowth, as measured 6h after puparation. See quantification in H, experimental design in I and representative images in J. Scale bars represent 50 μm. Wilcoxon rank-sum statistic test for two samples was performed in D, I and L.. * P<0.05, ** P<0.01 *** P<0.001. N.S. No statistical difference.

**Sup. Fig.2 related to Figure 2. Validation of a minimal 20E responsive reporter**

**A**. Expression of a dominant negative form of EcR inhibits the 10xERE-NLS4xNeonGreeen reporter in wing imaginal discs. Expression is induced in the dorsal compartment with *ap-Gal4*, leaving the ventral compartment as control. **B**. Quantification of whole live larva fluorescence produced by the reporter around the time of pupariation. Each line represents the average NeonGreen signal for a single pupa. Dashed line marks head eversion (h.e.). **C-D**. Activation of the 10xERE-NLS4xNG reporter in transgenic explanted imaginal discs treated with 20nM 20E. Time after addition of 20E is indicated. Robust activation is seen after 2.5h. The quantification for several discs (n=3) is shown in B and representative images are shown in D. Error bars represent standard deviation. Scale bars represent 50 μm.

**Sup. Fig.3 related to Figure 3. Role of EcR in disc eversion**.

A. High concentration (200 or 2000nM) but not low concentration (20nM) of 20E triggers eversion (recognised as tissue elongation) in wild type mid 3^rd^ instar imaginal discs after 18h of incubation. This does not occur if EcR is deleted from the pouch ΔEcR; EcR-GFP^cKO^ pdm2-Gal4 UAS-Flp).

**Sup. Fig.4 related to Figure 4. Expression of known 20E-regulated genes in explanted discs treated with various 20E concentrations**.

**A**. Normalised reading counts for genes known to be affected (directly or indirectly) by 20E. **B-E** Levels of Br-Z1-GFP and Blimp1-GFP, as assessed with anti-GFP, rise with increasing levels of 20E in explanted discs. **F**. For the upregulated genes, the canonical EcR binding site is the second most abundant TF binding motif present in the vicinity of the gene. **G**. This is not the case for downregulated genes. **H**. Example of read counts for genes from each of the three clusters. The Pearson correlation coefficient between the fold change and the δ_(2000−20)_/δ_(20−0)_ ratio was calculated for the experimental data (611 genes up-regulated by ecdysone) (**I**) and for data randomly generated (see Material and Methods) (**J**).The two correlation coefficients differed statistically (Fisher’s z-Tests for differences of correlations in two independent samples z = 8.28 with probability = 0).

**Sup. Fig.5 related to Figure 4. Models of gene regulation by Ecdysone**.

**A**. Diagram representing the molecular complexes that EcR (blue) makes with 20E (purple dot), its coactivator (green), and corepressor (red). Arrows indicate reversible binding events with affinity specified by the ratio of *on* and *off* rates). We also consider a truncated form of EcR lacking a DNA binding domain (EcD_LBD_), which acts as a “ligand sponge” to sequester ecdysone and other EcR binders. **B**. Probability for any given EcR to be found in each of the four accessible complexes described in panel A as a function of 20E and EcR concentration. The concentration of corepressor is assumed constant. As expected, EcR is mostly found in its activating form (EcRea) when EcR levels are low and 20E level are sufficiently high to expel the corepressor. Similarly, the repressive form of EcR (EcRr) is predominant in the low EcR, low 20E range. Parameters: κ_*A*_ = 1 nM^−1^, κ_*E*_ = 0.017; nM^−1^, κ_*R*_ = 0.017; nM^−1^, A_*tot*_ = 5 nM, R_*tot*_ = 30 nM, 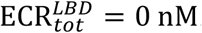. **C**. Visualisation of the context-dependent transcriptional action of a functional ERE, obtained by assigning a “modulation factor” to each EcR complex and computing a statistical average drawing on the probabilities of the ERE being occupied by EcR *and* the EcR being in a particular complex (see Mathematical Modeling in Material and Methods). Left: When the sponge is not expressed 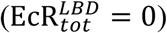, transcriptional activity is impaired by removing or overexpressing the EcR. Right: effect of various concentrations of EcD_LBD_ with EcR fixed at ECR_*tot*_ = 4 nM. Other parameters: κ_*A*_ = 1 nM^−1^, κ_*E*_ = 0.017; nM^−1^, κ_*R*_ = 0.017; nM^−1^, A_*tot*_ = 5 nM, R_*tot*_ = 30 nM, κ_*D*_ = 20 nM^−1^, ERE_*tot*_ = 1 nM. At low 20E, increasing EcD_LBD_ leads to derepression, while at high 20E levels it causes deactivation, recapitulating the results of ^61^. **D**. Probability of a set of EcRs effectively acting as transcriptional activator 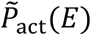, transcriptional inhihibor 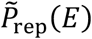 or a neutral unliganded element 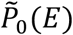 in three different models: left, one regulating EcR; middle, two regulating EcRs, right, 10 regulating EcRs. See Supplementary Informations for details and additional assumptions. Parameters: 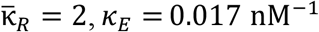. **E**. Effect of enhancer and silencer elements on the activity of ERE. Addition of a silencer (orange curve) decreases the overall response of ERE (purple and red curves), while an enhancer (lime curve) enforces increased baseline activity (purple and green curves). Equivalently, the ERE modulates the enhancer’s baseline activity by 20E-dependent switching from repressor to activator (lime and green curves). Silencers or enhancers modulate the concentration (and hence developmental time) at which high and low target genes cross a hypothetical threshold (black dashed and dotted curves, respectively), thus mimicking the mechanism that leads to nested expression of morphogen target genes. **F**. Fitting of the reporter activity as a function of ecdysone for different constructs (data shown in Fig. 6A-C) with the thermodynamic model using the two alternative expressions the probabilities 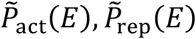 and 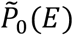 plotted in panel D (middle and right). Cooperative effects among the EREs lead to a sharper overall response of gene expression as a function as ecdysone concentration compared to the model with only ERE (Fig. 6D), which is in better agreement with the experimental data.

**Sup. Fig.6 related to Figure 5. Expression of the four synthetic reporters in freshly explanted discs**.

**A**. 2xBrkS-10xERE-NLS4xNG is only detectably expressed after 48h AL2-L3, when ecdysone levels have sufficiently risen. Fluorescence at 0 and 24 hrs originates from the trachea, which cannot be readily dissected away at these stages. **B**. As expected, the control construct (with mutated EREs) is not expressed. C. 3xGBE-10xERE-NLS By comparing **C** and **D**, we can infer EcR’s activity: At 0h, 44h and 48h AL2-L3 it acts as an activator, as the signal is higher in C than D. Whereas at 24h AL2-L3 it inhibits the transcription of the reporter. The activity at 0h might be due to the 20E peak that triggers L2 to L3 transition.

**Sup. Fig.7 related to Figure 6. Reporter expression data used for cross experiment comparison**.

**A**. Expression of 10xERE-NLS4xNG and 3xGBE-10xERE-NLS4xNG after 2.5 hr culture in 200nM 20E. At this concentration, 3xGBE-10xERE-NLS4xNG is expressed 1.4 times higher than NLS4xNG. This information was used to allow comparison of the quantifications displayed in Fig. 6 which came from images taken at different times.

**Sup. Vid. 1 Related to Sup.Fig. 2. Fluorescence from the 10xERE-NLS4xNG reporter during pupal stages**.

The line around the pupae shows the region where the average intensity of the signal was measured. The dynamic expression of the reporter followed the 20E titers. Scale bar represents 1 mm.

**Sup. Vid. 2 Related to Fig. 3A Cell proliferation in wing discs cultured ex-vivo without the addition of 20E**.

**Sup. Vid. 3 Related to Fig. 3A Cell proliferation in wing discs cultured ex-vivo with 10nM of 20E**

**Sup. Vid. 4 Related to Fig. 3A Cell proliferation in wing discs cultured ex-vivo with 20nM of 20E**

**Sup. Vid. 5 Related to Fig. 3A Cell proliferation in wing discs cultured ex-vivo with 40nM of 20E**

**Sup. Vid. 6 Related to Fig. 3D Lack of cell proliferation in wing discs cultured ex-vivo with 2000nM of 20E**

**Sup. Vid. 7 Related to Fig. 3B Cell proliferation in wing discs cultured ex-vivo without the addition of 20E**

**Sup. Vid. 8 Related to Fig. 3B Cell proliferation in wing discs cultured ex-vivo with 20nM of 20E**

**Sup. Vid. 9 Related to Fig. 3B Cell proliferation in wing discs lacking EcR and cultured ex-vivo without the addition of 20E**

Raw video is on the left and the region of interest (ROI) on the right. E-Cadherin is in green, and His-2B in red. Mitoses were manually marked with a blue dot in the ROI. Refer to Fig. 3 for more details. Scale bar represents 50 μm.

## Notes

### Competing Interest Statement

The authors have declared no competing interest.

## REFERENCES

1. Petkovich, M., and Chambon, P. (2022). Retinoic acid receptors at 35 years. J Mol Endocrinol 69, T13–T24. 10.1530/JME-22-0097.

2. Cunningham, T.J., and Duester, G. (2015). Mechanisms of retinoic acid signalling and its roles in organ and limb development. Nat Rev Mol Cell Biol 16, 110–123. 10.1038/nrm3932.

3. Niederreither, K., and Dolle, P. (2008). Retinoic acid in development: towards an integrated view. Nat Rev Genet 9, 541–553. 10.1038/nrg2340.

4. Brent, G.A. (2012). Mechanisms of thyroid hormone action. J Clin Invest 122, 3035–3043. 10.1172/JCI60047.

5. Liu, Y.C., Yeh, C.T., and Lin, K.H. (2019). Molecular Functions of Thyroid Hormone Signaling in Regulation of Cancer Progression and Anti-Apoptosis. Int J Mol Sci 20. 10.3390/ijms20204986.

6. Amoutzias, G.D., Pichler, E.E., Mian, N., De Graaf, D., Imsiridou, A., Robinson-Rechavi, M., Bornberg-Bauer, E., Robertson, D.L., and Oliver, S.G. (2007). A protein interaction atlas for the nuclear receptors: properties and quality of a hub-based dimerisation network. BMC Syst Biol 1, 34. 10.1186/1752-0509-1-34.

7. Frigo, D.E., Bondesson, M., and Williams, C. (2021). Nuclear receptors: from molecular mechanisms to therapeutics. Essays Biochem 65, 847–856. 10.1042/EBC20210020.

8. Mishra, S., Kelly, K.K., Rumian, N.L., and Siegenthaler, J.A. (2018). Retinoic Acid Is Required for Neural Stem and Progenitor Cell Proliferation in the Adult Hippocampus. Stem Cell Reports 10, 1705–1720. 10.1016/j.stemcr.2018.04.024.

9. Mosher, K.I., and Schaffer, D.V. (2018). Proliferation versus Differentiation: Redefining Retinoic Acid’s Role. Stem Cell Reports 10, 1673–1675. 10.1016/j.stemcr.2018.05.011.

10. Kress, E., Samarut, J., and Plateroti, M. (2009). Thyroid hormones and the control of cell proliferation or cell differentiation: paradox or duality? Mol Cell Endocrinol 313, 36–49. 10.1016/j.mce.2009.08.028.

11. Mouillet, J.F., Henrich, V.C., Lezzi, M., and Vogtli, M. (2001). Differential control of gene activity by isoforms A, B1 and B2 of the Drosophila ecdysone receptor. Eur J Biochem 268, 1811–1819.

12. Hu, X., Cherbas, L., and Cherbas, P. (2003). Transcription activation by the ecdysone receptor (EcR/USP): identification of activation functions. Mol Endocrinol 17, 716–731. 10.1210/me.2002-0287.

13. Dobens, L., Rudolph, K., and Berger, E.M. (1991). Ecdysterone regulatory elements function as both transcriptional activators and repressors. Mol Cell Biol 11, 1846–1853. 10.1128/mcb.11.4.1846.

14. Cherbas, L., Hu, X., Zhimulev, I., Belyaeva, E., and Cherbas, P. (2003). EcR isoforms in Drosophila: testing tissue-specific requirements by targeted blockade and rescue. Development 130, 271–284.

15. Tsai, C.C., Kao, H.Y., Yao, T.P., McKeown, M., and Evans, R.M. (1999). SMRTER, a Drosophila nuclear receptor coregulator, reveals that EcR-mediated repression is critical for development. Mol Cell 4, 175–186.

16. Bai, J., Uehara, Y., and Montell, D.J. (2000). Regulation of invasive cell behavior by taiman, a Drosophila protein related to AIB1, a steroid receptor coactivator amplified in breast cancer. Cell 103, 1047–1058.

17. Francis, V.A., Zorzano, A., and Teleman, A.A. (2010). dDOR is an EcR coactivator that forms a feed-forward loop connecting insulin and ecdysone signaling. Curr Biol 20, 1799–1808. 10.1016/j.cub.2010.08.055.

18. Schulman, I.G., Juguilon, H., and Evans, R.M. (1996). Activation and repression by nuclear hormone receptors: hormone modulates an equilibrium between active and repressive states. Mol Cell Biol 16, 3807–3813. 10.1128/MCB.16.7.3807.

19. Glass, C.K., and Rosenfeld, M.G. (2000). The coregulator exchange in transcriptional functions of nuclear receptors. Genes & development 14, 121–141.

20. Boulan, L., and Leopold, P. (2021). What determines organ size during development and regeneration? Development 148. 10.1242/dev.196063.

21. Texada, M.J., Koyama, T., and Rewitz, K. (2020). Regulation of Body Size and Growth Control. Genetics 216, 269–313. 10.1534/genetics.120.303095.

22. Pan, X., Connacher, R.P., and O’Connor, M.B. (2021). Control of the insect metamorphic transition by ecdysteroid production and secretion. Curr Opin Insect Sci 43, 11–20. 10.1016/j.cois.2020.09.004.

23. Kannangara, J.R., Mirth, C.K., and Warr, C.G. (2021). Regulation of ecdysone production in Drosophila by neuropeptides and peptide hormones. Open Biol 11, 200373. 10.1098/rsob.200373.

24. Kamiyama, T., and Niwa, R. (2022). Transcriptional Regulators of Ecdysteroid Biosynthetic Enzymes and Their Roles in Insect Development. Front Physiol 13, 823418. 10.3389/fphys.2022.823418.

25. Petryk, A., Warren, J.T., Marques, G., Jarcho, M.P., Gilbert, L.I., Kahler, J., Parvy, J.P., Li, Y., Dauphin-Villemant, C., and O’Connor, M.B. (2003). Shade is the Drosophila P450 enzyme that mediates the hydroxylation of ecdysone to the steroid insect molting hormone 20-hydroxyecdysone. Proc Natl Acad Sci U S A 100, 13773–13778. 10.1073/pnas.2336088100.

26. Okamoto, N., Viswanatha, R., Bittar, R., Li, Z., Haga-Yamanaka, S., Perrimon, N., and Yamanaka, N. (2018). A Membrane Transporter Is Required for Steroid Hormone Uptake in Drosophila. Dev Cell 47, 294–305 e297. 10.1016/j.devcel.2018.09.012.

27. Strassburger, K., Lorbeer, F.K., Lutz, M., Graf, F., Boutros, M., and Teleman, A.A. (2017). Oxygenation and adenosine deaminase support growth and proliferation of ex vivo cultured Drosophila wing imaginal discs. Development 144, 2529–2538. 10.1242/dev.147538.

28. Strassburger, K., Lutz, M., Muller, S., and Teleman, A.A. (2021). Ecdysone regulates Drosophila wing disc size via a TORC1 dependent mechanism. Nat Commun 12, 6684. 10.1038/s41467-021-26780-0.

29. Herboso, L., Oliveira, M.M., Talamillo, A., Perez, C., Gonzalez, M., Martin, D., Sutherland, J.D., Shingleton, A.W., Mirth, C.K., and Barrio, R. (2015). Ecdysone promotes growth of imaginal discs through the regulation of Thor in D. melanogaster. Sci Rep 5, 12383. 10.1038/srep12383.

30. Dye, N.A., Popovic, M., Spannl, S., Etournay, R., Kainmuller, D., Ghosh, S., Myers, E.W., Julicher, F., and Eaton, S. (2017). Cell dynamics underlying oriented growth of the Drosophila wing imaginal disc. Development 144, 4406–4421. 10.1242/dev.155069.

31. Parker, J., and Struhl, G. (2020). Control of Drosophila wing size by morphogen range and hormonal gating. Proc Natl Acad Sci U S A 117, 31935–31944. 10.1073/pnas.2018196117.

32. Nogueira Alves, A., Oliveira, M.M., Koyama, T., Shingleton, A., and Mirth, C.K. (2022). Ecdysone coordinates plastic growth with robust pattern in the developing wing. Elife 11. 10.7554/eLife.72666.

33. Guo, Y., Flegel, K., Kumar, J., McKay, D.J., and Buttitta, L.A. (2016). Ecdysone signaling induces two phases of cell cycle exit in Drosophila cells. Biol Open 5, 1648–1661. 10.1242/bio.017525.

34. O’Keefe, D.D., Thomas, S.R., Bolin, K., Griggs, E., Edgar, B.A., and Buttitta, L.A. (2012). Combinatorial control of temporal gene expression in the Drosophila wing by enhancers and core promoters. BMC Genomics 13, 498. 10.1186/1471-2164-13-498.

35. Schubiger, M., Carre, C., Antoniewski, C., and Truman, J.W. (2005). Ligand-dependent de-repression via EcR/USP acts as a gate to coordinate the differentiation of sensory neurons in the Drosophila wing. Development 132, 5239–5248. 10.1242/dev.02093.

36. Mirth, C.K., Truman, J.W., and Riddiford, L.M. (2009). The ecdysone receptor controls the post-critical weight switch to nutrition-independent differentiation in Drosophila wing imaginal discs. Development 136, 2345–2353. 10.1242/dev.032672.

37. Frame, K.K., and Hu, W.S. (1990). Cell volume measurement as an estimation of mammalian cell biomass. Biotechnol Bioeng 36, 191–197. 10.1002/bit.260360211.

38. Hecht, S., Perez-Mockus, G., Schienstock, D., Recasens-Alvarez, C., Merino-Aceituno, S., Smith, M., Salbreux, G., Degond, P., and Vincent, J.P. (2022). Mechanical constraints to cell-cycle progression in a pseudostratified epithelium. Curr Biol 32, 2076–2083 e2072. 10.1016/j.cub.2022.03.004.

39. Fain, M.J., and Stevens, B. (1982). Alterations in the cell cycle of Drosophila imaginal disc cells precede metamorphosis. Dev Biol 92, 247–258. 10.1016/0012-1606(82)90169-5.

40. Bryant, P.J., and Simpson, P. (1984). Intrinsic and extrinsic control of growth in developing organs. Q Rev Biol 59, 387–415.

41. Neufeld, T.P., de la Cruz, A.F., Johnston, L.A., and Edgar, B.A. (1998). Coordination of growth and cell division in the Drosophila wing. Cell 93, 1183–1193.

42. Martin, F.A., Herrera, S.C., and Morata, G. (2009). Cell competition, growth and size control in the Drosophila wing imaginal disc. Development 136, 3747–3756. 10.1242/dev.038406.

43. Worley, M.I., Setiawan, L., and Hariharan, I.K. (2013). TIE-DYE: a combinatorial marking system to visualize and genetically manipulate clones during development in Drosophila melanogaster. Development 140, 3275–3284. 10.1242/dev.096057.

44. Blanco-Obregon, D., El Marzkioui, K., Brutscher, F., Kapoor, V., Valzania, L., Andersen, D.S., Colombani, J., Narasimha, S., McCusker, D., Leopold, P., and Boulan, L. (2022). A Dilp8-dependent time window ensures tissue size adjustment in Drosophila. Nat Commun 13, 5629. 10.1038/s41467-022-33387-6.

45. Ohhara, Y., Kobayashi, S., and Yamanaka, N. (2017). Nutrient-Dependent Endocycling in Steroidogenic Tissue Dictates Timing of Metamorphosis in Drosophila melanogaster. PLoS Genet 13, e1006583. 10.1371/journal.pgen.1006583.

46. Talamillo, A., Herboso, L., Pirone, L., Perez, C., Gonzalez, M., Sanchez, J., Mayor, U., Lopitz-Otsoa, F., Rodriguez, M.S., Sutherland, J.D., and Barrio, R. (2013). Scavenger receptors mediate the role of SUMO and Ftz-f1 in Drosophila steroidogenesis. PLoS Genet 9, e1003473. 10.1371/journal.pgen.1003473.

47. Gibbens, Y.Y., Warren, J.T., Gilbert, L.I., and O’Connor, M.B. (2011). Neuroendocrine regulation of Drosophila metamorphosis requires TGFbeta/Activin signaling. Development 138, 2693–2703. 10.1242/dev.063412.

48. Uyehara, C.M., and McKay, D.J. (2019). Direct and widespread role for the nuclear receptor EcR in mediating the response to ecdysone in Drosophila. Proc Natl Acad Sci U S A. 10.1073/pnas.1900343116.

49. Koelle, M.R., Talbot, W.S., Segraves, W.A., Bender, M.T., Cherbas, P., and Hogness, D.S. (1991). The Drosophila EcR gene encodes an ecdysone receptor, a new member of the steroid receptor superfamily. Cell 67, 59–77.

50. Milner, M.J. (1977). The eversion and differentiation of Drosophila melanogaster leg and wing imaginal discs cultured in vitro with an optimal concentration of β-ecdysone. Development 37, 105–117.

51. Martin, P., and Shearn, A. (1980). Development of Drosophila imaginal discs in vitro: effects of ecdysone concentration and insulin. Journal of Experimental Zoology 211, 291–301.

52. Lavrynenko, O., Rodenfels, J., Carvalho, M., Dye, N.A., Lafont, R., Eaton, S., and Shevchenko, A. (2015). The ecdysteroidome of Drosophila: influence of diet and development. Development 142, 3758–3768. 10.1242/dev.124982.

53. Shlyueva, D., Stelzer, C., Gerlach, D., Yanez-Cuna, J.O., Rath, M., Boryn, L.M., Arnold, C.D., and Stark, A. (2014). Hormone-responsive enhancer-activity maps reveal predictive motifs, indirect repression, and targeting of closed chromatin. Mol Cell 54, 180–192. 10.1016/j.molcel.2014.02.026.

54. Akagi, K., and Ueda, H. (2011). Regulatory mechanisms of ecdysone-inducible Blimp-1 encoding a transcriptional repressor that is important for the prepupal development in Drosophila. Dev Growth Differ 53, 697–703. 10.1111/j.1440-169X.2011.01276.x.

55. Shea, M.A., and Ackers, G.K. (1985). The OR control system of bacteriophage lambda. A physical-chemical model for gene regulation. J Mol Biol 181, 211–230. 10.1016/0022-2836(85)90086-5.

56. Buchler, N.E., Gerland, U., and Hwa, T. (2003). On schemes of combinatorial transcription logic. Proc Natl Acad Sci U S A 100, 5136–5141. 10.1073/pnas.0930314100.

57. Parker, D.S., White, M.A., Ramos, A.I., Cohen, B.A., and Barolo, S. (2011). The cis-regulatory logic of Hedgehog gradient responses: key roles for gli binding affinity, competition, and cooperativity. Sci Signal 4, ra38. 10.1126/scisignal.2002077.

58. Sherman, M.S., and Cohen, B.A. (2012). Thermodynamic state ensemble models of cis-regulation. PLoS Comput Biol 8, e1002407. 10.1371/journal.pcbi.1002407.

59. Cohen, M., Page, K.M., Perez-Carrasco, R., Barnes, C.P., and Briscoe, J. (2014). A theoretical framework for the regulation of Shh morphogen-controlled gene expression. Development 141, 3868–3878. 10.1242/dev.112573.

60. Bintu, L., Buchler, N.E., Garcia, H.G., Gerland, U., Hwa, T., Kondev, J., and Phillips, R. (2005). Transcriptional regulation by the numbers: models. Curr Opin Genet Dev 15, 116–124. 10.1016/j.gde.2005.02.007.

61. Wardwell-Ozgo, J., Terry, D., Schweibenz, C., Tu, M., Solimon, O., Schofeld, D., and Moberg, K. (2022). An EcR probe reveals mechanisms of the ecdysone-mediated switch from repression-to-activation on target genes in the larval wing disc. bioRxiv, 2022.2004.2007.487542. 10.1101/2022.04.07.487542.

62. Muller, B., Hartmann, B., Pyrowolakis, G., Affolter, M., and Basler, K. (2003). Conversion of an extracellular Dpp/BMP morphogen gradient into an inverse transcriptional gradient. Cell 113, 221–233. 10.1016/s0092-8674(03)00241-1.

63. Furriols, M., and Bray, S. (2001). A model Notch response element detects Suppressor of Hairless-dependent molecular switch. Curr Biol 11, 60–64. 10.1016/s0960-9822(00)00044-0.

64. Junker, J.P., Peterson, K.A., Nishi, Y., Mao, J., McMahon, A.P., and van Oudenaarden, A. (2014). A predictive model of bifunctional transcription factor signaling during embryonic tissue patterning. Dev Cell 31, 448–460. 10.1016/j.devcel.2014.10.017.

65. Delas, M.J., and Briscoe, J. (2020). Repressive interactions in gene regulatory networks: When you have no other choice. Curr Top Dev Biol 139, 239–266. 10.1016/bs.ctdb.2020.03.003.

66. Cavallo, R.A., Cox, R.T., Moline, M.M., Roose, J., Polevoy, G.A., Clevers, H., Peifer, M., and Bejsovec, A. (1998). Drosophila Tcf and Groucho interact to repress Wingless signalling activity. Nature 395, 604–608. 10.1038/26982.

67. van de Wetering, M., Cavallo, R., Dooijes, D., van Beest, M., van Es, J., Loureiro, J., Ypma, A., Hursh, D., Jones, T., Bejsovec, A., et al. (1997). Armadillo coactivates transcription driven by the product of the Drosophila segment polarity gene dTCF. Cell 88, 789–799. 10.1016/s0092-8674(00)81925-x.

68. Andres, A.J., Fletcher, J.C., Karim, F.D., and Thummel, C.S. (1993). Molecular analysis of the initiation of insect metamorphosis: a comparative study of Drosophila ecdysteroid-regulated transcription. Dev Biol 160, 388–404. 10.1006/dbio.1993.1315.

69. Zhang, C., Robinson, B.S., Xu, W., Yang, L., Yao, B., Zhao, H., Byun, P.K., Jin, P., Veraksa, A., and Moberg, K.H. (2015). The ecdysone receptor coactivator Taiman links Yorkie to transcriptional control of germline stem cell factors in somatic tissue. Dev Cell 34, 168–180. 10.1016/j.devcel.2015.05.010.

70. Wang, C., Yin, M.X., Wu, W., Dong, L., Wang, S., Lu, Y., Xu, J., Wu, W., Li, S., Zhao, Y., and Zhang, L. (2016). Taiman acts as a coactivator of Yorkie in the Hippo pathway to promote tissue growth and intestinal regeneration. Cell Discov 2, 16006. 10.1038/celldisc.2016.6.

71. Uyehara, C.M., Leatham-Jensen, M., and McKay, D.J. (2022). Opportunistic binding of EcR to open chromatin drives tissue-specific developmental responses. Proc Natl Acad Sci U S A 119, e2208935119. 10.1073/pnas.2208935119.

72. Russell, S.R., Heimbeck, G., Goddard, C.M., Carpenter, A.T., and Ashburner, M. (1996). The Drosophila Eip78C gene is not vital but has a role in regulating chromosome puffs. Genetics 144, 159–170. 10.1093/genetics/144.1.159.

73. Schenk, T., Stengel, S., and Zelent, A. (2014). Unlocking the potential of retinoic acid in anticancer therapy. Br J Cancer 111, 2039–2045. 10.1038/bjc.2014.412.

74. Tang, X.H., and Gudas, L.J. (2011). Retinoids, retinoic acid receptors, and cancer. Annu Rev Pathol 6, 345–364. 10.1146/annurev-pathol-011110-130303.

75. Krashin, E., Piekielko-Witkowska, A., Ellis, M., and Ashur-Fabian, O. (2019). Thyroid Hormones and Cancer: A Comprehensive Review of Preclinical and Clinical Studies. Front Endocrinol (Lausanne) 10, 59. 10.3389/fendo.2019.00059.

76. Moeller, L.C., and Fuhrer, D. (2013). Thyroid hormone, thyroid hormone receptors, and cancer: a clinical perspective. Endocr Relat Cancer 20, R19–29. 10.1530/ERC-12-0219.

77. Dye, N.A. (2022). Cultivation and Live Imaging of Drosophila Imaginal Discs. Methods Mol Biol 2540, 317–334. 10.1007/978-1-0716-2541-5_16.

78. Kim, D., Paggi, J.M., Park, C., Bennett, C., and Salzberg, S.L. (2019). Graph-based genome alignment and genotyping with HISAT2 and HISAT-genotype. Nat Biotechnol 37, 907–915. 10.1038/s41587-019-0201-4.

79. Li, H., Handsaker, B., Wysoker, A., Fennell, T., Ruan, J., Homer, N., Marth, G., Abecasis, G., Durbin, R., and Genome Project Data Processing, S. (2009). The Sequence Alignment/Map format and SAMtools. Bioinformatics 25, 2078–2079. 10.1093/bioinformatics/btp352.

80. Liao, Y., Smyth, G.K., and Shi, W. (2014). featureCounts: an efficient general purpose program for assigning sequence reads to genomic features. Bioinformatics 30, 923–930. 10.1093/bioinformatics/btt656.

81. Yoo, A.B., Jette, M.A., and Grondona, M. (2003). SLURM: Simple Linux Utility for Resource Management. held in Berlin, Heidelberg, (Springer Berlin Heidelberg), pp. 44–60.

82. Love, M.I., Huber, W., and Anders, S. (2014). Moderated estimation of fold change and dispersion for RNA-seq data with DESeq2. Genome Biol 15, 550. 10.1186/s13059-014-0550-8.

83. Mächler, M., Rousseeuw, P., Struyf, A., Hubert, M., and Hornik, K. (2012). Cluster: Cluster Analysis Basics and Extensions.

84. Shannon, P., Markiel, A., Ozier, O., Baliga, N.S., Wang, J.T., Ramage, D., Amin, N., Schwikowski, B., and Ideker, T. (2003). Cytoscape: a software environment for integrated models of biomolecular interaction networks. Genome Res 13, 2498–2504. 10.1101/gr.1239303.

85. Verfaillie, A., Imrichova, H., Janky, R., and Aerts, S. (2015). iRegulon and i-cisTarget: Reconstructing Regulatory Networks Using Motif and Track Enrichment. Curr Protoc Bioinformatics 52, 2 16 11–12 16 39. 10.1002/0471250953.bi0216s52.

86. Janky, R., Verfaillie, A., Imrichova, H., Van de Sande, B., Standaert, L., Christiaens, V., Hulselmans, G., Herten, K., Naval Sanchez, M., Potier, D., et al. (2014). iRegulon: from a gene list to a gene regulatory network using large motif and track collections. PLoS Comput Biol 10, e1003731. 10.1371/journal.pcbi.1003731.

