## Supplementary material for "The ecdysone receptor promotes or suppresses proliferation according to ligand level": Mathematical Modelling

February 8, 2023

In this supplementary text, we present in Section 1 a simple reaction model describing the equilibrium statistics of the ecdysone (20E) nuclear receptor (EcR) binding to different EcR ligands as well as an ecdysone responsive element (ERE), its DNA binding site. Our mathematical framework recapitulates recent experimental results [8] in which expression of a diffusible EcR fragment lacking a DNA binding domain was observed to produce opposite effects on reporter transcription, depending on 20E level. In Section 2, we describe the thermodynamic model of transcription used to inform the design of the reporters incorporating basal enhancers and silencers, which we refer to collectively as *basal regulatory modules* in the following. We point out generic features of this family of models and suggest how they can be refined to incorporate higher order cooperative interactions between repeated elements, particularly EREs, in order to more closely recapitulate both physiological and synthetic 20E target responses.

### 1 Statistics of EcR complex formation

In this section we discuss an equilibrium model for the occupancy of the ecdysone responsive element (ERE) by the ecdysone nuclear receptor (EcR). In the first section "General model and effect of EcR ligand sponge" we derive the ERE occupancy probabilities by different EcR complexes, taking into account possible roles of the co-repressor and co-activator of the EcR, as well as the presence of 20E-binding fragments $\mathrm{EcR}^{\mathrm{LBD}}$. These fragments can bind the co-repressor and co-activator but do not bind to the ERE, and are introduced here for comparison with the experiments of [8]. In the second section "Chemostatting of free EcR ligands and large binding affinity of EcR to ERE leads to Hill-type response of ecdysone-regulated genes" we obtain the ERE occupancy probabilities under simplifying assumptions and a situation without 20E-binding fragments $\mathrm{EcR}^{\mathrm{LBD}}$.

#### 1.1 General model and effect of EcR ligand sponge

In the following, we denote EcR and $\mathrm{EcR}^{\mathrm{LBD}}$ the concentration of 20E receptors and 20E-binding fragments, respectively, that are not bound to any ligand and not bound to an ERE. $\mathrm{EcR}^{\mathrm{LBD}}$, which acts as an EcR ligand "sponge", is assumed to lack a DNA binding domain, therefore not to be able to bind to the ERE and to be found only in its free, diffusible form. We denote $\mathrm{EcR}_{\mathrm{r}}$, the complex of EcR and the co-repressor Smarter (Smr), $\mathrm{EcR}_{\mathrm{e}}$, the complex of EcR and 20E, and $\mathrm{EcR}_{\mathrm{e,a}}$, the complex of EcR, 20E and the putative co-activator. The presence of such a co-activation is not essential for the model to exhibit the features of interest, however we allow for its existence here for the sake of generality (further simplifications are introduced in Section 1.2). Similarly, we denote $\mathrm{EcR}_{\mathrm{r}}^{\mathrm{LBD}}$, $\mathrm{EcR}_{\mathrm{e}}^{\mathrm{LBD}}$ and $\mathrm{EcR}_{\mathrm{e,a}}^{\mathrm{LBD}}$ the complexes of $\mathrm{EcR}^{\mathrm{LBD}}$ with the co-repressor, 20E, and 20E and the co-activator respectively. The concentrations of free binders are denoted $R$, $A$ and E for the co-repressor, co-activator and 20E, respectively. We refer the reader to Sup. Fig. 5A for a schematic representation of the reaction network.

As we consider an equilibrium reaction model, at steady state the mean-field reaction equations are solved by the detailed balance (zero flux) conditions

$$k_{\text{off}}^R \text{EcR}_{\text{r}}^{(\text{LBD})} = k_{\text{on}}^R \times R \times \text{EcR}^{(\text{LBD})} \tag{S1}$$

$$k_{\text{off}}^E \text{EcR}_{\text{e}}^{(\text{LBD})} = k_{\text{on}}^E \times \text{E} \times \text{EcR}^{(\text{LBD})} \tag{S2}$$

$$k_{\text{off}}^A \text{EcR}_{\text{e,a}}^{(\text{LBD})} = k_{\text{on}}^A \times A \times \text{EcR}_{\text{e}}^{(\text{LBD})} \tag{S3}$$

where parenthesis indicate that the equation holds for EcR or $\text{EcR}^{\text{LBD}}$; together with the expressions for the conservation of the total receptor concentration:

$$\text{EcR}_{\text{tot}} = \text{EcR} + \text{EcR}_{\text{r}} + \text{EcR}_{\text{e}} + \text{EcR}_{\text{e,a}} \tag{S4}$$

$$\text{EcR}_{\text{tot}}^{\text{LBD}} = \text{EcR}^{\text{LBD}} + \text{EcR}_{\text{r}}^{\text{LBD}} + \text{EcR}_{\text{e}}^{\text{LBD}} + \text{EcR}_{\text{e,a}}^{\text{LBD}} \tag{S5}$$

as well as the total ligand concentration:

$$R = R_{\text{tot}} - \text{EcR}_{\text{r}} - \text{EcR}_{\text{r}}^{\text{LBD}} \tag{S6}$$

$$\text{E} = \text{E}_{\text{tot}} - \text{EcR}_{\text{e}} - \text{EcR}_{\text{e}}^{\text{LBD}} - \text{EcR}_{\text{e,a}} - \text{EcR}_{\text{e,a}}^{\text{LBD}} \tag{S7}$$

$$A = A_{\text{tot}} - \text{EcR}_{\text{e,a}} - \text{EcR}_{\text{e,a}}^{\text{LBD}} \ , \tag{S8}$$

where the subscript $\bullet_{\text{tot}}$ refers to the total concentration of a given species, independent of whether it appears as part of a complex.

Eqs. (S1)-(S8) can be used to obtain the steady-state probabilities of all species. We want to determine the steady-state probabilities of a given EcR being found in one of its complexes:

$$P(\text{EcR}_{\text{e}}) = \frac{\text{EcR}_{\text{e}}}{\text{EcR}_{\text{tot}}} \tag{S9}$$

$$P(\text{EcR}_{\text{e,a}}) = \frac{\text{EcR}_{\text{e,a}}}{\text{EcR}_{\text{tot}}} \tag{S10}$$

$$P(\text{EcR}_{\text{r}}) = \frac{\text{EcR}_{\text{r}}}{\text{EcR}_{\text{tot}}} \ , \tag{S11}$$

as we later assume that these probabilities then modulate transcription of the 20E-regulated genes. Having defined the ligand binding affinities

$$\kappa_R = \frac{k_{\text{on}}^R}{k_{\text{off}}^R}, \quad \kappa_E = \frac{k_{\text{on}}^E}{k_{\text{off}}^E}, \quad \kappa_A = \frac{k_{\text{on}}^A}{k_{\text{off}}^A} \tag{S12}$$

it is now convenient to express the steady-state concentrations of the different receptor complexes in terms of EcR and $\text{EcR}^{\text{LBD}}$ only:

$$\begin{aligned} \text{EcR}_{\text{r}}^{(\text{LBD})} &= \kappa_R \times R \times \text{EcR}^{(\text{LBD})} \\ \text{EcR}_{\text{e}}^{(\text{LBD})} &= \kappa_E \times \text{E} \times \text{EcR}^{(\text{LBD})} \\ \text{EcR}_{\text{e,a}}^{(\text{LBD})} &= (\kappa_A A)(\kappa_E \text{E}) \times \text{EcR}^{(\text{LBD})} \end{aligned} \tag{S13}$$

with

$$R = \frac{R_{\text{tot}}}{1 + \kappa_R(\text{EcR} + \text{EcR}^{\text{LBD}})} \tag{S14}$$

$$A = \frac{A_{\text{tot}}}{1 + \kappa_A \kappa_E \text{E}(\text{EcR} + \text{EcR}^{\text{LBD}})} \tag{S15}$$

$$\text{E} = \frac{\text{E}_{\text{tot}}}{1 + \kappa_E(1 + \kappa_A A_f)(\text{EcR} + \text{EcR}^{\text{LBD}})} \ . \tag{S16}$$

Using Eq. (S4) we can further write

$$\text{EcR} = \frac{\text{EcR}_{\text{tot}}}{1 + \kappa_R R + \kappa_E \text{E} + \kappa_A \kappa_E A \text{E}} \ . \tag{S17}$$

The ligand binding properties of the nuclear receptor EcR and its fragment $\mathrm{EcR}^{\mathrm{LBD}}$ are assumed to be identical. Therefore, the relative concentrations of the various complexes involving the EcR are not affected by the absence of the DNA binding domain, notably:

$$\frac{\mathrm{EcR}}{\mathrm{EcR}_{\mathrm{tot}}} = \frac{\mathrm{EcR}^{\mathrm{LBD}}}{\mathrm{EcR}^{\mathrm{LBD}}_{\mathrm{tot}}} \,, \tag{S18}$$

and thus

$$\mathrm{EcR} + \mathrm{EcR}^{\mathrm{LBD}} = \mathrm{EcR}\left(1 + \frac{\mathrm{EcR}^{\mathrm{LBD}}_{\mathrm{tot}}}{\mathrm{EcR}_{\mathrm{tot}}}\right) \,, \tag{S19}$$

which can be used to eliminate $\mathrm{EcR}^{\mathrm{LBD}}$ for the equations above. To make further progress we would therefore like to solve Eq. (S17) for EcR as a function of $\mathrm{EcR}_{\mathrm{tot}}$. The concentrations of free ligands ($A_f$, $R_f$, $\mathrm{E}_f$) themselves depend on EcR via Eqs. (S14)-(S16). Eqs. (S15) and (S16) need to be dealt with first, since they are coupled. After some algebra we find

$$A = \frac{-\mathcal{K} + \sqrt{\mathcal{K}^2 + 4\kappa_E\kappa_A A_{\mathrm{tot}}(\mathrm{EcR} + \mathrm{EcR}^{\mathrm{LBD}})\left(1 + \kappa_E(\mathrm{EcR} + \mathrm{EcR}^{\mathrm{LBD}})\right)}}{2\kappa_E\kappa_A(\mathrm{EcR} + \mathrm{EcR}^{\mathrm{LBD}})} \tag{S20}$$

with

$$\mathcal{K} = 1 + \kappa_E(\mathrm{EcR} + \mathrm{EcR}^{\mathrm{LBD}})(1 + \kappa_A(\mathrm{E}_{\mathrm{tot}} - A_{\mathrm{tot}})) \,. \tag{S21}$$

and Eq. (S16) then yields E as a function of EcR, $\mathrm{EcR}^{\mathrm{LBD}}$. We can now determine $P(\mathrm{EcR_e})$, $P(\mathrm{EcR_{e,a}})$ and $P(\mathrm{EcR_r})$. This is done by substituting Eqs. (S14), (S20) and (S16) into Eq. (S17), which is subsequently solved for EcR numerically in Wolfram Mathematica, and finally using Eq. (S13). The results are shown in Sup.Fig. 5B.

In order to regulate the transcription of 20E targets, the ecdysone nuclear receptor (EcR) needs to be bound to DNA at the ecdysone responsive element (ERE). Assuming that the affinity of the EcR for the ERE, $\kappa_D$, is independent of the specific EcR complex, the probability $P_b$ that a given ERE is occupied by an EcR is the positive solution of

$$P_b = \frac{\kappa_D(\mathrm{EcR}_{\mathrm{tot}} - P_b\mathrm{ERE}_{\mathrm{tot}})}{1 + \kappa_D(\mathrm{EcR}_{\mathrm{tot}} - P_b\mathrm{ERE}_{\mathrm{tot}})} \tag{S22}$$

where $\mathrm{ERE}_{\mathrm{tot}}$ denotes the total concentration of ERE (bound or unbound to an EcR) and $\mathrm{EcR}_{\mathrm{tot}} - P_b\mathrm{ERE}_{\mathrm{tot}} \equiv \mathrm{EcR}_f$ is the concentration of free nuclear receptors. Note that, by assumption, this probability does not depend on the concentration of the EcR ligands. The probability that a given ERE is occupied by a specific EcR complex is thus obtained by multiplying any of the above mentioned $P(\mathrm{EcR}_\bullet)$ by the binding probability $P_b$.

Assuming that both the unoccupied ERE and the unliganded form of EcR have no net transcriptional effect, it follows that 20E levels can only results in a modulation of the trancriptional activity of a target gene if the total receptor concentration $\mathrm{EcR}_{\mathrm{tot}}$ is comparable to $\mathrm{ERE}_{\mathrm{tot}}$. This is due to the fact that at low $\mathrm{EcR}_{\mathrm{tot}}$ the binding probability $P_b$ is low and EREs are mostly unoccupied, while a progressive increase in $\mathrm{EcR}_{\mathrm{tot}}$ at fixed total ligands concentration leads to depletion of the latters and thus a low probability for EcR to be found in its ligated form (Sup. Fig. 5C).

To visualise net transcriptional regulatory action predicted by the model as a function of its free parameters and at a qualitative level, we assign a “modulation factor” $m$ to each EcR complex and compute a statistical average drawing on the steady-state probabilities determined above. For the sake of simplicity we assign a factor $m = +1$ (indicating activation) to the DNA-bound $\mathrm{EcR_{e,a}}$ complex, a factor $m = -1$ (indicating repression) to the DNA-bound $\mathrm{EcR_r}$ complex and we let $m = 0$ (indicating neutral action) for all other configurations. The average is thus given by

$$\langle m\rangle = P_b \times (P(\mathrm{EcR_{e,a}}) - P(\mathrm{EcR_r})) \tag{S23}$$

and plotted in Sup. Fig. 5C for the case of $\mathrm{EcR}^{\mathrm{LBD}}_{\mathrm{tot}} = 0$ (left) and as a function of $\mathrm{EcR}^{\mathrm{LBD}}_{\mathrm{tot}}$ at fixed $\mathrm{EcR}_{\mathrm{tot}} = 4\mathrm{nM}$ (right). In agreement with the experimental results of [8], we observe that $\mathrm{EcR}^{\mathrm{LBD}}$ expression leads to de-repression at low 20E and de-activation at high 20E levels.

#### 1.2 Chemostatting of free EcR ligands and large binding affinity of EcR to ERE leads to Hill-type response of 20E-regulated genes

For the purpose of this study, we do not need to consider the role of 20E binding fragments, so that we now assume that $\mathrm{EcR}_{\mathrm{tot}}^{\mathrm{LBD}} = 0$. A quantitative comparison to our results, as well as analytical insight, is made impractical due to the large number of unknown parameters. To make progress, we rely on simplifying assumptions. In the following, we impose that (a) the concentration of free EcR ligands ($\mathrm{E} = \mathrm{E}_{\mathrm{tot}}, A = A_{\mathrm{tot}}, R = R_{\mathrm{tot}}$) is chemostatted, i.e. constant and thus not subject to depletion effects; (b) the concentration of EcR is large compared to that of the EREs ($\mathrm{EcR}_{\mathrm{tot}} \gg \mathrm{ERE}_{\mathrm{tot}}$), (c) binding between EcR and ERE happens with high affinity ($\kappa_D \gg 1$), such that $P_b \simeq 1$, i.e. we can assume that any given ERE is at all times occupied by an EcR, (d) the role of transcriptional co-activators can be neglected, so that $A_f = 0$.

We then obtain the following probabilities that a given ERE is occupied by a particular EcR complex:

$$P_{\mathrm{rep}}(\mathrm{E}) \equiv P(\mathrm{EcR_r}) = \frac{\mathrm{EcR_r}}{\mathrm{EcR_{tot}}} = \frac{\bar{\kappa}_R}{1 + \bar{\kappa}_R + \kappa_E \mathrm{E}} \tag{S24}$$

$$P_0(\mathrm{E}) \equiv P(\mathrm{EcR}) = \frac{\mathrm{EcR}}{\mathrm{EcR_{tot}}} = \frac{1}{1 + \bar{\kappa}_R + \kappa_E \mathrm{E}} \tag{S25}$$

$$P_{\mathrm{act}}(\mathrm{E}) \equiv P(\mathrm{EcR_e}) = \frac{\mathrm{EcR_e}}{\mathrm{EcR_{tot}}} = \frac{\kappa_E \mathrm{E}}{1 + \bar{\kappa}_R + \kappa_E \mathrm{E}}\ , \tag{S26}$$

such that $P_{\mathrm{rep}}(\mathrm{E}) + P_0(\mathrm{E}) + P_{\mathrm{act}}(\mathrm{E}) = 1$ and where the only unknown parameter is the effective affinity of the co-repressor, $\bar{\kappa}_R = \kappa_R R$, while $\kappa_E \simeq 1.7 \times 10^{-2}\ \mathrm{nM}^{-1}$ [7].

Here, we use the notation $P_{\mathrm{act}}(\mathrm{E})$ to refer to the probability that a given 20E-regulated gene has an ERE bound to an EcR in its active state, as a function of the concentration of free 20E, E. We recognise (S26) as a Hill function of order $h = 1$ (Michaelis-Menten) with characteristic concentration $\mathrm{E}^0 = (1 + \bar{\kappa}_R)/\kappa_E$. This is plotted in Sup. Fig. 5D. $P_{\mathrm{rep}}(\mathrm{E})$ and $P_0(\mathrm{E})$ correspond to the probability that a given ERE has an EcR bound in a repressive or unliganded state.

### 2 Thermodynamic model of transcription regulation by EcR

In this section, we introduce the thermodynamic model of transcriptional regulation by the ecdysone nuclear receptor EcR that was used to inform the design of the synthetic reporters presented in the main text. This class of mathematical models is well established in the literature [5, 2, 1, 4, 6, 3] and connects the occupancy of the promoter region by different transcription factors (TF), here the EcR and additional factors recruited by the basal regulatory modules, to the binding of the basal transcriptional complex (BTC) to the promoter and the resulting average transcriptional rate (see schematic in Fig. 4F). We distinguish two classes for these additional factors: transcriptional activators which can bind to enhancers, and transcriptional repressors which can bind to silencers. We distinguish five types of regulatory architectures, which we label ERE, Enh, Sil, Enh-ERE and Sil-ERE; where Enh, Sil, ERE in the name indicates the presence of an enhancer, silencer and ERE respectively.

#### 2.1 Single ERE site

We start our analysis by considering a single ERE site, possibly in combination with a basal regulatory module. For the sake of simplicity, we assume here that transcriptional activators and repressors are permanently bound to enhancers and silencers, and that all ERE sites are occupied by an EcR. In the absence of bound enhancers, silencers, and for unliganded EcR, the effective affinity of the BTC for the promoter is denoted $\kappa_p$. This affinity is modulated by cooperativity coefficients depending on the state of enhancer, silencers and DNA-bound EcR. The affinity of the BTC for the promoter $\kappa_p$ is thus multiplied by the positive factors $C_{\mathrm{TA}}$, $C_{\mathrm{TR}}$, $C_{\mathrm{EA}}$ and $C_{\mathrm{ER}}$ in the presence of a transcriptional activator, transcriptional repressor, EcR in an activating state (bound to 20E) and EcR in a repressive state (bound to a corepressor), respectively. As a result of these definitions, we expect $C_{\rm TA} > 1$, $C_{\rm TR} < 1$, $C_{\rm EA} > 1$ and $C_{\rm ER} < 1$. In the thermodynamic model of transcription, the cooperativity coefficients $C_{\rm TA}$, $C_{\rm TR}$, $C_{\rm EA}$ and $C_{\rm ER}$ reflect the differences in free energy of the configurations where the BTC is bound to the promoter, in the presence or absence of transcriptional activators and repressors.

Expressing the probability of the BTC to be bound to the promoter, we then obtain the following expressions for the transcriptional activity of different configurations:

$$A_{\rm ERE}({\rm E}) = k_T\kappa_p\frac{P_0({\rm E}) + C_{\rm ER}P_{\rm rep}({\rm E}) + C_{\rm EA}P_{\rm act}({\rm E})}{1+\kappa_p(P_0({\rm E}) + C_{\rm ER}P_{\rm rep}({\rm E}) + C_{\rm EA}P_{\rm act}({\rm E}))}\ , \tag{S27}$$

$$A_{\rm Enh}({\rm E}) = k_T\frac{\kappa_p C_{\rm TA}}{1+\kappa_p C_{\rm TA}}\ , \tag{S28}$$

$$A_{\rm Sil}({\rm E}) = k_T\frac{\kappa_p C_{\rm TR}}{1+\kappa_p C_{\rm TR}}\ , \tag{S29}$$

$$A_{\rm Enh-ERE}({\rm E}) = k_T\kappa_p C_{\rm TA}\frac{P_0({\rm E}) + C_{\rm ER}P_{\rm rep}({\rm E}) + C_{\rm EA}P_{\rm act}({\rm E})}{1+\kappa_p C_{\rm TA}(P_0({\rm E}) + C_{\rm ER}P_{\rm rep}({\rm E}) + C_{\rm EA}P_{\rm act}({\rm E}))}\ , \tag{S30}$$

$$A_{\rm Sil-ERE}({\rm E}) = k_T\kappa_p C_{\rm TR}\frac{P_0({\rm E}) + C_{\rm ER}P_{\rm rep}({\rm E}) + C_{\rm EA}P_{\rm act}({\rm E})}{1+\kappa_p C_{\rm TR}(P_0({\rm E}) + C_{\rm ER}P_{\rm rep}({\rm E}) + C_{\rm EA}P_{\rm act}({\rm E}))}\ , \tag{S31}$$

where the probabilities $P_0$, $P_{\rm rep}$, $P_{\rm act}$ are defined in Eqs. (S24)-(S26), and $k_T$ corresponds to the rate of gene expression, when the BTC is bound to the promoter. Note that in this context $P_0$, $P_{\rm rep}$ and $P_{\rm act}$ no longer correspond to the probability for the ERE to be occupied by a particular EcR complex. This is due to the reciprocity of the cooperative interactions, which can equivalently be seen as stabilising or destabilising (depending on the value of the corresponding coefficient) the binding of the transcriptional co-factors to the EcR in the presence of a bound BTC, thus affecting the occupation statistics (see Eq. (S38) below).

To analyse these expressions (S27)-(S31) for the transcriptional activity of the different configurations, we focus on the general form:

$$A({\rm E}) = k_T\kappa_p C_{\rm T}\frac{P_0({\rm E}) + C_{\rm ER}P_{\rm rep}({\rm E}) + C_{\rm EA}P_{\rm act}({\rm E})}{1+\kappa_p C_{\rm T}(P_0({\rm E}) + C_{\rm ER}P_{\rm rep}({\rm E}) + C_{\rm EA}P_{\rm act}({\rm E}))} \tag{S32}$$

which reduces to a pure ERE for $C_{\rm T} = 1$, to a pure enhancer for $C_{\rm ER} = C_{\rm EA} = 1$ and $C_{\rm T} > 1$, to a pure silencer for $C_{\rm ER} = C_{\rm EA} = 1$ and $C_{\rm T} < 1$, and describes the behaviour of a basal silencer combined with an ERE for $C_{\rm T} < 1$ and a basal enhancer combined with an ERE for $C_{\rm T} > 1$. Therefore, the dependency of $A({\rm E})$ on $C_{\rm T}$ allows to evaluate the effect on transcriptional regulation of adding enhancers and silencers of different strengths. We now discuss these effects.

First, we obtain the following expression for the fold-change $f = \lim_{{\rm E}\to\infty} A({\rm E})/A(0)$:

$$f = \frac{C_{\rm EA}(C_{\rm T}\kappa_p(1+C_{\rm ER}\bar{\kappa}_R) + \bar{\kappa}_R + 1)}{(1+C_{\rm ER}\bar{\kappa}_R)(1+C_{\rm EA}C_{\rm T}\kappa_p)}\ , \tag{S33}$$

which decreases, as $C_{\rm T}$ spans the range from 0 to $\infty$, from $f = C_{\rm EA}(1+\bar{\kappa}_R)/(1+C_{\rm ER}\bar{\kappa}_R)$ to $f = 1$. The fold change $f$ can be seen as the dynamic range of gene expression as 20E is varied, and is maximal for a perfect basal silencer $C_{\rm T} = 0$ and minimal for a perfect basal enhancer, $C_{\rm T} \to \infty$ (using that $C_{\rm ER} < 1$, $C_{\rm EA} > 1$). Using that $C_{\rm ER} < 1$, $C_{\rm EA} > 1$, one can also verify that the fold change is maximal for small $\kappa_p$ and large values of $C_{EA}$; corresponding to a gene with a low basal transcriptional activity which is strongly enhanced by 20E. Fitting of reporter data suggests that the EcR/ERE is indeed in that regime (Section 2.4).

By first calculating the difference between the transcriptional activity $A$ at a given concentration of 20E and its value at zero 20E concentration, and subsequently normalising to the new maximum, we defined a scaled version of the transcriptional activity:

$$\tilde{A}({\rm E}) = \frac{A({\rm E}) - A(0)}{A({\rm E}\to\infty) - A(0)}\ . \tag{S34}$$

Defining the threshold activation value of 20E as the 20E concentration at which $\tilde{A} = \frac{1}{2}$, we obtain:

$$\mathrm{E}^* = \frac{1}{\kappa_E} \frac{1 + \bar{\kappa}_R + C_\mathrm{T}\kappa_p(1 + C_\mathrm{ER}\bar{\kappa}_R)}{1 + C_\mathrm{EA}C_\mathrm{T}\kappa_p} \,. \tag{S35}$$

Here as well, the threshold activation value of 20E decreases with $C_T$, from $\mathrm{E}^* = (1 + \bar{\kappa}_R)/\kappa_E$ at $C_\mathrm{T} = 0$ to $(1 + C_\mathrm{ER}\kappa_R)/(C_\mathrm{EA}\kappa_E)$ for $C_\mathrm{T} \to \infty$, where we again assume $C_\mathrm{EA} > 1$ and $C_\mathrm{ER} < 1$.

Therefore, increasing the strength of a basal enhancer (as measured by $C_\mathrm{T}$) leads to early activation of a gene by 20E, as well as a lower dynamic range, as measured by the fold change $f$. The ratio between $f$ and $E^*$ is independent of $C_\mathrm{T}$ and is given by:

$$\frac{f}{\mathrm{E}^*} = \frac{C_\mathrm{EA}\kappa_E}{1 + C_\mathrm{ER}\bar{\kappa}_R} \,. \tag{S36}$$

To experimentally test this relationship, we use the following ratio obtained from experimental data (Figs. 4E, 4H), instead of the threshold concentration $E^*$:

$$\frac{\delta_{(2000-20)}}{\delta_{(20-0)}} \equiv \frac{A(2000\ \mathrm{nM}) - A(20\ \mathrm{nM})}{A(20\ \mathrm{nM}) - A(0\ \mathrm{nM})}. \tag{S37}$$

which gives an indirect estimate of the threshold activation value of 20E; as for simple, monotonous activation curves, genes which reach their maximal activation at concentration larger than 20nM ($E^* > 20$nM) would also tend to have larger ratios of $\delta_{(2000-20)}/\delta_{(20-0)}$.

Finally, as we hinted at earlier, we note that the probability of an EcR bound to ERE to be also bound to 20E is different from the probability of a free EcR to be bound to 20E, $P_\mathrm{act}(\mathrm{E})$, and reads:

$$P_{\mathrm{ERE-EcR-E}}(\mathrm{E}) = \frac{\tilde{\kappa}_E \mathrm{E}}{1 + \bar{\kappa}_R + \tilde{\kappa}_E\mathrm{E}} = \frac{\mathrm{E}}{\mathrm{E} + \mathrm{E}^*}, \quad \tilde{\kappa}_E = \kappa_E \frac{(1 + \bar{\kappa}_R)(1 + \kappa_p C_T C_\mathrm{EA})}{1 + \bar{\kappa}_R + \kappa_p C_T(1 + C_\mathrm{ER}\bar{\kappa}_R)} \,. \tag{S38}$$

Eq. (S38) shows that the dependence of the activation threshold of the EcR by 20E, E$^*$, on the various cooperativity coefficients established in Eq. (S35), can equivalently be understood in terms of a renormalisation of the affinity of 20E for a ERE-bound EcR, $\tilde{\kappa}_E$, by the cooperative interactions.

This renormalised affinity is larger than the bare affinity $\kappa_E$ when $C_\mathrm{EA} > 1 + (C_\mathrm{ER} - 1)\bar{\kappa}_R/(1 + \bar{\kappa}_R)$, which is always satisfied when $C_\mathrm{EA} > 1$ and $C_\mathrm{ER} < 1$, as we assume here. Indeed, the fact that the affinity for the promoter of the BTC increases when the EcR is bound to 20E implies in the thermodynamic model that the EcR-E is stabilised in return when the BTC is bound. Because the presence of a basal enhancer favors BTC binding to the promoter, the thermodynamic model predicts that this also leads to increased affinity of 20E for the EcRs bound to an ERE: $\tilde{\kappa}_E$ increases with $C_\mathrm{T}$ provided that $C_\mathrm{EA} > 1$ and $C_\mathrm{ER} < 1$. We note that this implies that the characterisation of the affinity of a nuclear receptor in isolation might miss important features originating from its interactions with the BTC, as well as other basal regulatory elements.

Finally, we can obtain the 20E concentration at which the EcR changes from acting as a repressor to acting as an activator for a given gene, at fixed $C_\mathrm{T}$. This is obtained by solving $A(\mathrm{E}) = A(\mathrm{E})|_{C_\mathrm{ER}=1, C_\mathrm{EA}=1}$ for the 20E concentration E$^{**}$; we obtain:

$$\mathrm{E}^{**} = \frac{\bar{\kappa}_R(1 - C_\mathrm{ER})}{\kappa_E(C_\mathrm{EA} - 1)}, \tag{S39}$$

which is independent of $C_\mathrm{T}$. With parameter values obtained by fitting experimental data (section 2.4), we obtain $\mathrm{E}^{**} \simeq 4.6$ nM. We note that models incorporating ERE cooperativity, describe below, leads to a shift of E$^{**}$ towards higher values (Sup. Fig. 5F).

#### 2.2 Cooperative action of two ERE sites

Reporters considered here in this work are expressed downstream of 10ERE sites, leaving the possibility of cooperative action between several bound EcRs. To explore the effect of cooperative action of EcRs, here we consider a model consisting of two EREs. As before, we consider that these ERE sites are permanently bound to an EcR. We assume that both EcRs must be bound to an 20E in order to positively modulate the binding affinity of the BTC for the promoter, with cooperativity coefficient $C_{\mathrm{EA}}$. Conversely, we assume that at least one of the two EcRs bound to a corepressor is sufficient to repress transcription, by reducing the binding affinity of the BTC for the promoter, with cooperativity coefficient $C_{\mathrm{ER}}$. We assume that all other states of EcR occupancy are neutral, in the sense that they do not affect the binding affinity of the BTC for the promoter. Basal transcriptional and regulators are treated as in the previous section. Calculating the transcriptional activity, we find that Eqs. (S27)-(S31) still apply (with the new definition for the coefficients $C_{\mathrm{EA}}$ and $C_{\mathrm{ER}}$, provided that the probabilities $P_{\mathrm{act}}(\mathrm{E})$, $P_{\mathrm{rep}}(\mathrm{E})$, $P_0(\mathrm{E})$ are replaced by the probabilities $\tilde{P}_{\mathrm{act}}(\mathrm{E})$, $\tilde{P}_{\mathrm{rep}}(\mathrm{E})$, $\tilde{P}_0(\mathrm{E})$, defined as follows:

$$\tilde{P}_{\mathrm{act}}(\mathrm{E}) = (P_{\mathrm{act}}(\mathrm{E}))^2 \tag{S40}$$

$$\tilde{P}_{\mathrm{rep}}(\mathrm{E}) = 1 - (1 - P_{\mathrm{rep}}(\mathrm{E}))^2 \tag{S41}$$

$$\tilde{P}_0(\mathrm{E}) = (1 - P_{\mathrm{rep}}(\mathrm{E}))^2 - (P_{\mathrm{act}}(\mathrm{E}))^2 \ . \tag{S42}$$

#### 2.3 Multiple ERE sites

We now consider a situation taking into account all of the 10 ERE sites of reporters considered in this study. We consider that all ERE sites are occupied by an EcR. We assume that at least 6 EcRs bound to 20E are required to positively modulate the binding affinity of the BTC for the promoter, with cooperativity coefficient $C_{\mathrm{EA}}$. We also assume that at least 5 EcR bound to a corepressor are required to repress BTC binding, with cooperativity coefficient $C_{\mathrm{ER}}$. Finally, we assume that all other states of occupancy of the bound EcRs are neutral and leave the effective binding affinity of the BTC to its basal value $\kappa_p$. Calculating the transcriptional activity, we find that Eqs. (S27)-(S31) still apply (with the new definition for the coefficients $C_{\mathrm{EA}}$ and $C_{\mathrm{ER}}$, provided that the probabilities $P_{\mathrm{act}}(E)$, $P_{\mathrm{rep}}(E)$, $P_0(E)$ are replaced by the probabilities $\tilde{P}_{\mathrm{act}}(E)$, $\tilde{P}_{\mathrm{rep}}(E)$, $\tilde{P}_0(E)$, defined as follows:

$$\tilde{P}_{\mathrm{act}}(\mathrm{E}) = \sum_{k=6}^{10} \binom{10}{k} (P_{\mathrm{act}}(\mathrm{E}))^k (1 - P_{\mathrm{act}}(\mathrm{E}))^{10-k} \tag{S43}$$

$$\tilde{P}_{\mathrm{rep}}(\mathrm{E}) = \sum_{k=5}^{10} \binom{10}{k} (P_{\mathrm{rep}}(\mathrm{E}))^k (1 - P_{\mathrm{rep}}(\mathrm{E}))^{10-k} \tag{S44}$$

$$\tilde{P}_0(\mathrm{E}) = 1 - \tilde{P}_{\mathrm{act}}(\mathrm{E}) - \tilde{P}_{\mathrm{rep}}(\mathrm{E}) \ . \tag{S45}$$

#### 2.4 Comparison with reporter data

We now describe the procedure that was followed to quantitatively compare the thermodynamic model described in the previous Section with the reporter signal data for the 10xERE-NLS4xNG, 2xBrkS-10xERE-NLS4xNG and 3XGBE-10XERE-NLS4xNG constructs, as well as for the control constructs 2xBrkS-10xERE*-NLS4xNG, 3XGBE-10XERE*-NLS4xNG with inactive EREs. The corresponding experimental data are shown in Fig. 6. For the sake of compactness, we refer to these constructs using the notation introduced in the previous section, ERE, Sil-ERE, Enh-ERE, Sil and Enh respectively.

The model equations (S27)-(S31), which assume that the dependence on 20E level of the collective action of the set of 10 EREs can be approximated by that of a single ERE, were fitted simultaneously to the five sets of reporter activity show in Fig. 6 A-C by minimising a loss function $L$ of the free parameters defined as the sum of the squared distances of the predicted response from the full set of experimental measurements,

$$L = \sum_{\mathrm{construct}} \sum_{E \in \{0,20,200,2000\}} \sum_{\mathrm{repeat}=1}^{7} \left[ A^{(\mathrm{model})}_{\mathrm{construct}}(\mathrm{E}) - A^{(\mathrm{exp})}_{\mathrm{construct}}(\mathrm{E}) \right]^2 \ . \tag{S46}$$

The minimisation was performed numerically in Python using the scipy.optimize package (Nelder-Mead method), returning the fitting parameters: $\kappa_p = 2.79 \times 10^{-2}, C_{\mathrm{TA}} = 1.91, C_{\mathrm{TR}} = 1.45 \times 10^{-1}, \kappa_R = 1.04, C_{\mathrm{ER}} = 8.75 \times 10^{-2}, C_{\mathrm{EA}} = 13.2$. Note that $C_{\mathrm{TR}}, C_{\mathrm{ER}} < 1$, while $C_{\mathrm{TA}}, C_{\mathrm{EA}} > 1$, in agreement with the assumed function of the corresponding elements, and in general showing weaker (positive or negative) cooperativity originating from the basal regulatory modules compared with the ERE sites, potentially due to the large number of repetitions of the latter. The initiation rate $k_T$ was also fitted to the reporter levels ($k_T = 1.31 \times 10^2$ a.u.) but its value can not be easily interpreted due to the units of fluorescence being arbitrary. The response curve predicted by the model after fitting is shown in Fig. 6D.

A similar procedure was followed to fit the form of the model where cooperative interactions between two or more ERE sites, which we described above. For the case of cooperative action between two ERE sites, Eqs. (S40)-(S42), we obtain the fitting parameters: $k_T = 60.3, \kappa_p = 3.94 \times 10^{-2}, C_{\mathrm{TA}} = 3.32, C_{\mathrm{TR}} = 8.72 \times 10^{-2}, \bar{\kappa}_R = 4.0 \times 10^{-1}, C_{\mathrm{ER}} = 5.24 \times 10^{-14}, C_{\mathrm{EA}} = 38.9$. On the other hand, for the case of 10 interacting ERE sites, Eqs. (S43)-(S45), we obtain the fitting parameters: $k_T = 47.6, \kappa_p = 6.49 \times 10^{-3}, C_{\mathrm{TA}} = 26.6, C_{\mathrm{TR}} = 7.42 \times 10^{-2}, \bar{\kappa}_R = 6.0 \times 10^{-1}, C_{\mathrm{ER}} = 6.96 \times 10^{-34}, C_{\mathrm{EA}} = 3.13 \times 10^2$. In both cases we find $C_{\mathrm{ER}} \simeq 0$, indicating a strong repressive action of the EcR-corepressor complex. The response curves predicted by the two variants of the model after fitting are shown in Sup. Fig. 5F. In both cases, the sharper activation between 20 nM and 200 nM appears to better match the experimental data (in fact, the value of the cost function (S46) is lowest across all three variations of the model at the minimum identified for the case of 10 interacting ERE sites).

### References


[1] Lacramioara Bintu, Nicolas E Buchler, Hernan G Garcia, Ulrich Gerland, Terence Hwa, Jané Kondev, and Rob Phillips. Transcriptional regulation by the numbers: models. *Current opinion in genetics & development*, 15(2):116–124, 2005.

[2] NE Buchler, U Gerland, and T Hwa. Predicting expression patterns from regulatory sequence in drosophila segmentation. *PNAS*, 100(9):5136–5141, 2003.

[3] Michael Cohen, Karen M Page, Ruben Perez-Carrasco, Chris P Barnes, and James Briscoe. A theoretical framework for the regulation of shh morphogen-controlled gene expression. *Development*, 141(20):3868–3878, 2014.

[4] David S Parker, Michael A White, Andrea I Ramos, Barak A Cohen, and Scott Barolo. The cis-regulatory logic of hedgehog gradient responses: key roles for gli binding affinity, competition, and cooperativity. *Science signaling*, 4(176):ra38–ra38, 2011.

[5] Madeline A Shea and Gary K Ackers. The or control system of bacteriophage lambda: A physical-chemical model for gene regulation. *Journal of molecular biology*, 181(2):211–230, 1985.

[6] Marc S Sherman and Barak A Cohen. Thermodynamic state ensemble models of cis-regulation. *PLoS computational biology*, 8(3):e1002407, 2012.

[7] S Tarlochan, TS Dhadialla, R Glenn, GR Carlson, and DP Le. New insecticides with ecdysteroidal and juvenile hormone activity. *Annu. Rev. Entomol*, 43:545–569, 1998.

[8] Joanna Wardwell-Ozgo, Douglas Terry, Colby Schweibenz, Michael Tu, Ola Solimon, David Schofeld, and Kenneth Moberg. An ecr probe reveals mechanisms of the ecdysone-mediated switch from repression-to-activation on target genes in the larval wing disc. *bioRxiv*, pages 2022–04, 2022.
