## Supplementary Figures for "The ecdysone receptor promotes or suppresses proliferation according to ligand level"

### Supplementary Figure 1

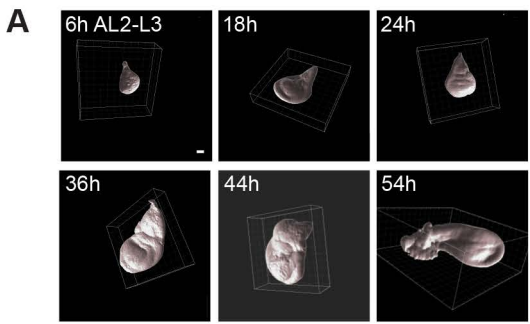

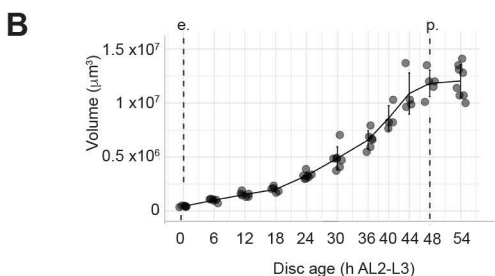

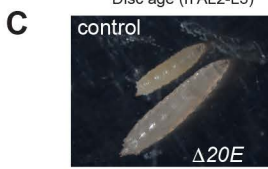

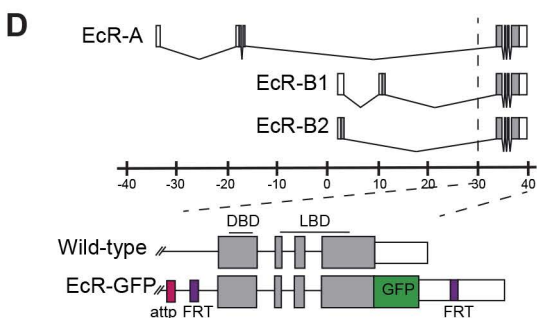

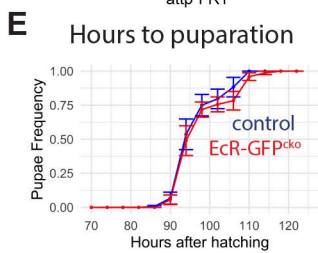

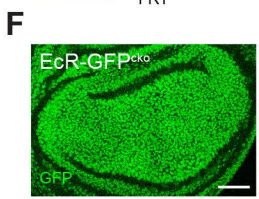

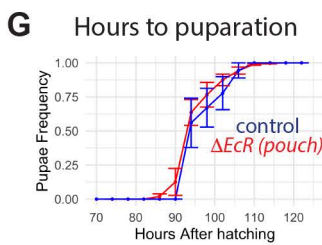

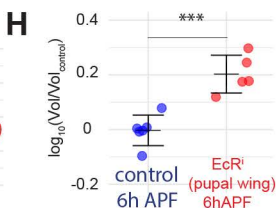

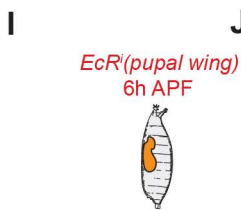

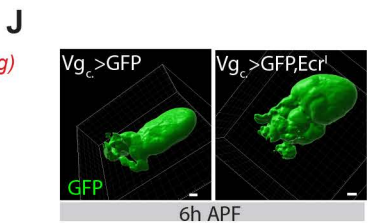

### Supplementary Figure 2

**A**

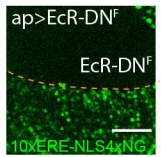

**B**

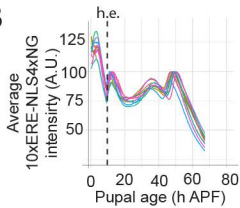

**C**

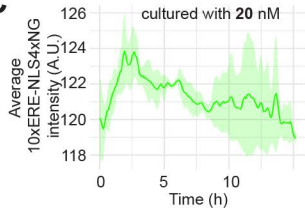

**D**

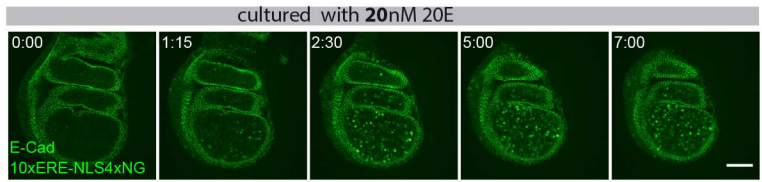

### Supplementary Figure 3

A

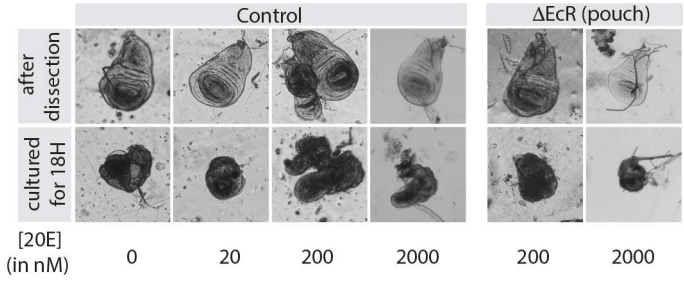

### Supplementary Figure 4

**A**

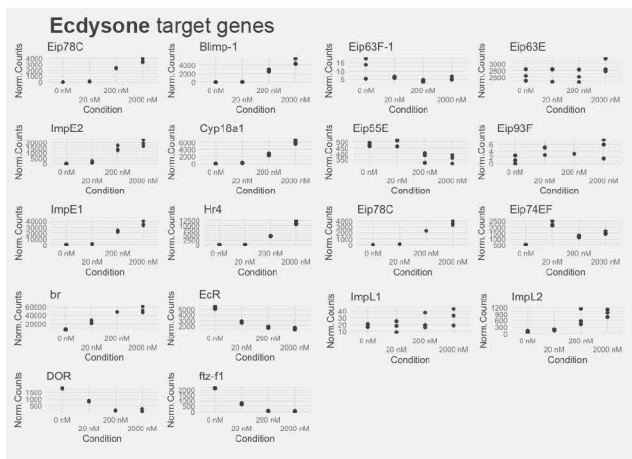

**B**

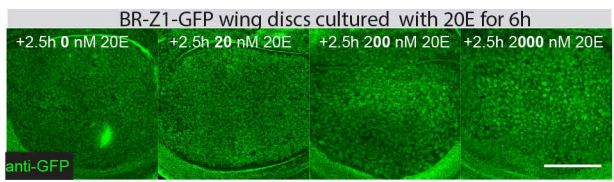

**C**

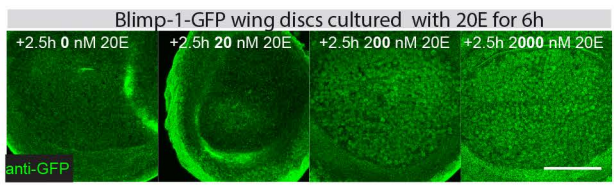

**D**

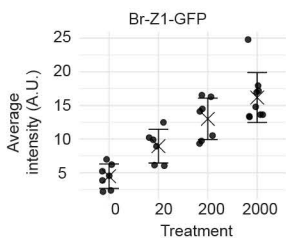

**E**

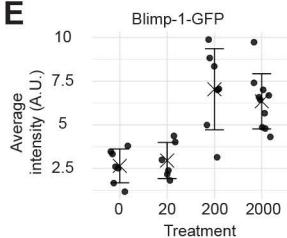

**F**

Top TF binding motifs enriched in **UPREGULATED** genes (607)

| Cluster | #TFs | TF | NES | #Targets | #Motifs |
| --- | --- | --- | --- | --- | --- |
| M1 | 1 | CG33260, Trl | 5.193 | 344 | 15 |
| M2 | 3 | EcR | 4.553 | 320 | 7 |
| M3 | 9 | ci, sug, lmd, opa | 4.502 | 157 | 11 |
| M4 | 2 | Med.pnr.pdm3.rn.br.cnc... | 4.266 | 384 | 30 |
| M5 | 10 | nau | 4.173 | 311 | 3 |

**G**

Top TF binding motifs enriched in **DOWNREGULATED** genes (635)

| Cluster | #TFs | TF | NES | #Targets | #Motifs |
| --- | --- | --- | --- | --- | --- |
| M1 | 2 | gem, Myb, zh1, grh | 4.807 | 300 | 9 |
| M2 | 6 | Jra, Atf3, Xbp1, CrebB-17A... | 4.679 | 305 | 33 |
| M3 | 10 | CG33260, Trl | 4.259 | 245 | 17 |
| M4 | 37 | tra2 | 3.872 | 244 | 3 |
| M5 | 10 | Top2 | 3.732 | 217 | 3 |

**H**

**Cluster 1**

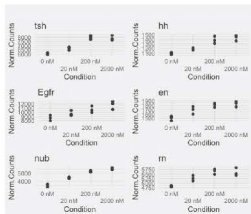

**Cluster 2**

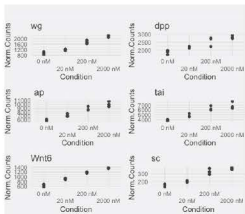

**Cluster 3**

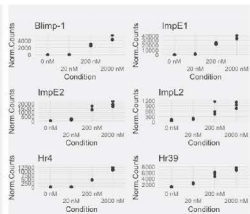

**I**

**Experimental data**

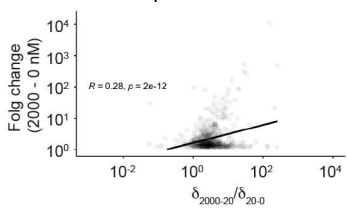

**J**

**Randomly generated data**

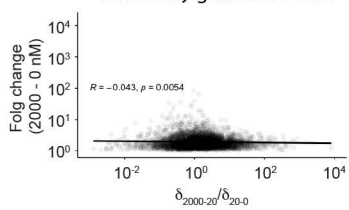

### Supplementary Figure 5

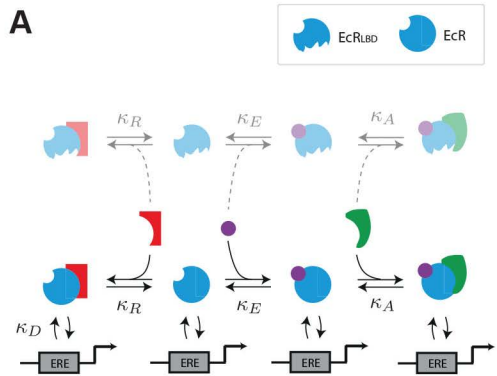

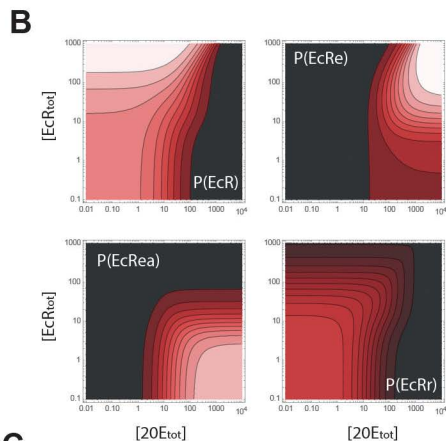

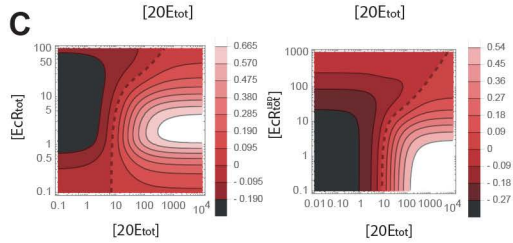

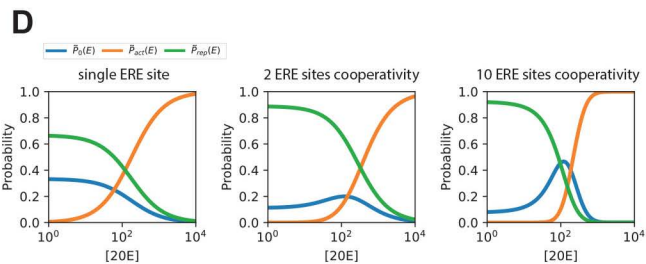

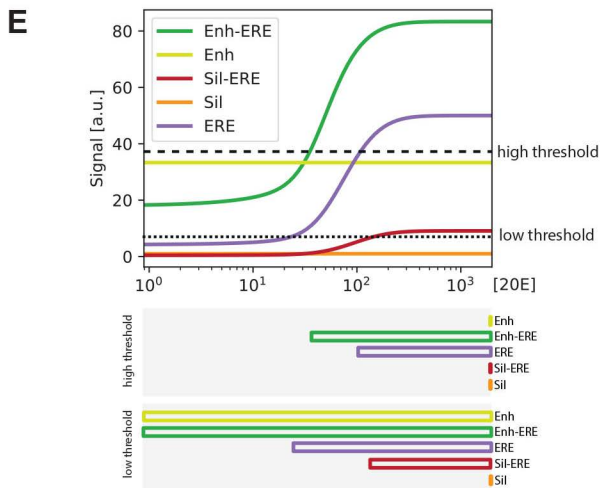

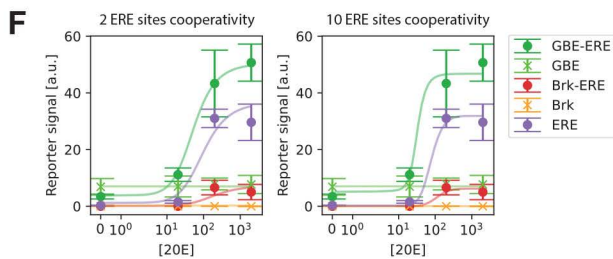

### Supplementary Figure 6

**A**

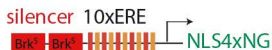

**2Brk<sup>5</sup>-10xERE-NLS4xNG**

non cultured

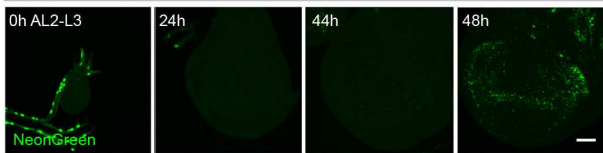

**B**

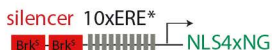

**2Brk<sup>5</sup>-10xERE\*-NLS4xNG**

non cultured

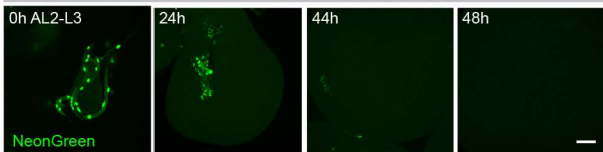

**C**

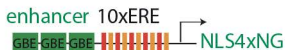

**3xGBE-10xERE-NLS4xNG**

non cultured

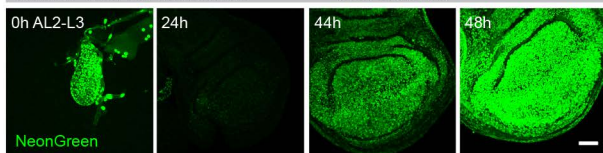

**D**

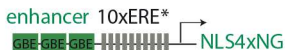

**3xGBE-10xERE\*-NLS4xNG**

non cultured

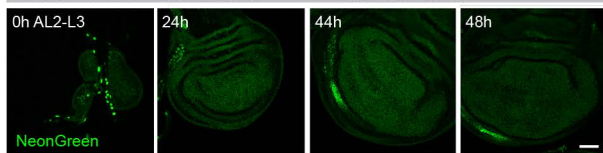

### Supplementary Figure 7

A

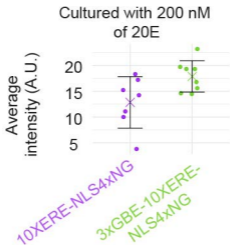
